# Metabolic bypass rescues aberrant S-nitrosylation-induced TCA cycle inhibition and synapse loss in Alzheimer’s disease human neurons

**DOI:** 10.1101/2023.10.12.562118

**Authors:** Alexander Y. Andreyev, Hongmei Yang, Paschalis-Thomas Doulias, Nima Dolatabadi, Xu Zhang, Melissa Luevanos, Mayra Blanco, Christine Baal, Ivan Putra, Tomohiro Nakamura, Harry Ischiropoulos, Steven R. Tannenbaum, Stuart A. Lipton

## Abstract

In Alzheimer’s disease (AD), dysfunctional mitochondrial metabolism is associated with synaptic loss, the major pathological correlate of cognitive decline. Mechanistic insight for this relationship, however, is still lacking. Here, comparing isogenic wild-type and AD mutant human induced pluripotent stem cell (hiPSC)-derived cerebrocortical neurons (hiN), we found evidence for compromised mitochondrial energy in AD using the Seahorse platform to analyze glycolysis and oxidative phosphorylation (OXPHOS). Isotope-labeled metabolic flux experiments revealed a major block in activity in the tricarboxylic acid (TCA) cycle at the α-ketoglutarate dehydrogenase (αKGDH)/succinyl coenzyme-A synthetase step, metabolizing α-ketoglutarate to succinate. Associated with this block we found aberrant protein S-nitrosylation of αKGDH subunits that are known to inhibit enzyme function. This aberrant S-nitrosylation was documented not only in AD-hiN but also in postmortem human AD brains vs. controls, as assessed by two separate unbiased mass spectrometry platforms using both *SNO*TRAP identification of S-nitrosothiols and chemoselective-enrichment of S-nitrosoproteins. Treatment with dimethyl succinate, a cell-permeable derivative of a TCA substrate (downstream to the block, resulted in partial rescue of mitochondrial bioenergetic function as well as reversal of synapse loss in AD-hiN. Our findings have therapeutic implications that rescue of mitochondrial energy metabolism can ameliorate synaptic loss in hiPSC-based models of AD.

## INTRODUCTION

Mitochondrial deficits and bioenergetic compromise have been proposed to contribute to neurodegenerative disorders, including Alzheimer’s disease (AD) (Gibson et al., 2010), and have been associated with synaptic loss in human AD brains assessed with specific PET markers (Venkataraman et al., 2022). Synaptic loss represents a critical neuropathological correlate to cognitive decline (Terry et al., 1991; DeKoskey et al., 1990; Tzioras et al., 2023). In part accounting for this energy loss, dysfunction of the mitochondrial tricarboxylic acid (TCA) cycle, involving particularly α-ketoglutarate dehydrogenase (αKGDH, also known as 2-oxoglutarate dehydrogenase), has been identified in several neurodegenerative disorders including AD (Gibson et al., 2010). Posttranslational modification of enzymes in the TCA cycle has been identified as an important nidus of control of bioenergetics, but little mechanistic work has been done in this area during disease processes (Arnold and Finley, 2022). Hence, the underlying mechanism(s) for TCA cycle dysfunction and the full extent of the energy compromise remain unknown. Accordingly, using human postmortem AD brain and hiPSCs to model AD, we report here the presence of aberrant protein S-nitrosylation of neuronal TCA cycle enzymes, including αKGDH subunits, which inhibits the activity of these enzymes, as we have previously shown (Doulias et al., 2021; Doulias et al., 2023). Recently, we reported the presence of ∼1,500 S-nitrosylation sites on proteins in AD brain and control brains (Yang et al., 2022). Among the aberrantly S-nitrosylated proteins in both male and female AD brains compared to controls, we noted that enzymes in metabolic pathways, particularly the TCA cycle, were affected (Yang et al., 2022). Here, using ^13^C dynamic labeling of AD hiPSC-derived cerebrocortical neurons (AD-hiN), we observed specific changes in central carbon metabolism indicating defects in the tricarboxylic acid (TCA) cycle that correlate to the enzymes manifesting aberrant protein S-nitrosylation. Additionally, using the Seahorse platform, we found that AD-hiN compared to isogenic WT/Controls manifested a decrease in maximal respiratory capacity (measured as oxygen consumption rate, OCR) but minimal effects basal glycolytic capacity (monitored by extracellular acidification rate, ECAR). These changes were also consistent with TCA cycle dysfunction in the AD-hiN. In accord with the AD-hiN findings, the highest level of aberrant S-nitrosylation in human postmortem brains with AD vs. controls was also found in subunits of αKGDH. Thus, our hiPSC-based model had pathophysiologically-relevant changes in S-nitrosylation of enzymes in the TCA cycle.

## RESULTS

### Protein S-nitrosylation of TCA enzymes in human AD brains and AD-hiN

Using a novel chemical probe specific for S-nitrosopeptides called *SNO*TRAP, we recently used mass spectrometry (MS) techniques for an assessment of the S-nitrosoproteome in 40 male and female human Alzheimer brains vs. control brains (Yang et al., 2022). Bioinformatic analysis of these data demonstrated that the TCA cycle enzymes were one of the highest scoring pathways affected by S-nitrosylation in both male and female AD brains vs. controls, and the #2 pathway in male AD brains (as shown in Table S12 and Figure S3 of Yang et al., 2022). By semiquantitative analysis of the SNO-proteins using spectral counting (Yang et al., 2022), the most upregulated SNO-TCA enzymes were aconitase (Aco, also known as aconitate hydratase, p = 0.021 by ANOVA) and dihydrolipoyl dehydrogenase (DLD, representing the E3 subunit of α-ketoglutarate dehydrogenase (αKGDH, also known as 2-oxoglutarate dehydrogenase) as well as of other dehydrogenases), p = 0.009 by ANOVA) (**Figures 1A** and **1B**). Malate dehydrogenase, mitochondrial (MDH), was somewhat more S-nitrosylated in AD brain than in controls (**Figure 1A**, p = 0.021 by ANOVA). Note, however, that malate dehydrogenase, cytoplasmic (also known as malic enzyme 2, ME2) (**Figure 1A**, p = 0.004 by ANOVA), was S-nitrosylated to an even greater degree in AD than control human brains, consistent with the notion that the alternative anaplerotic pathway access of malate or oxaloacetate into the TCA cycle may be regulated at this level by S-nitrosylation (**Figure 1B**).

**Figure 1.**
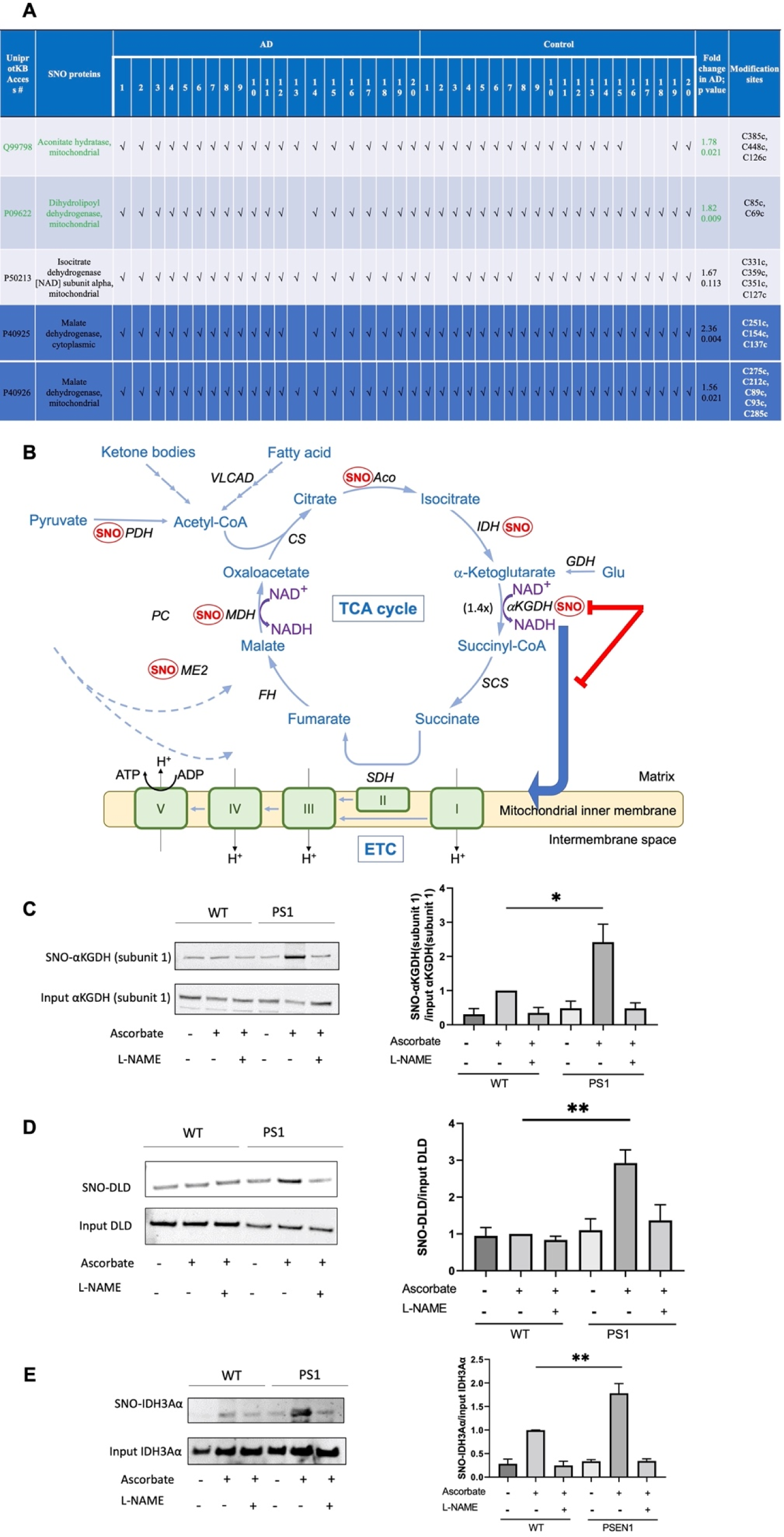
TCA enzymes significantly S-nitrosylated to a greater extent in AD brains than in control brains, as assessed by SNOTRAP/MS. (**A**) List of S-nitrosylated TCA enzymes in AD and Controls brains, assessed as in Yang et al. (2022). The first ten AD and Control brain samples are male, and second ten in each case are female. Fold increase in AD over Control shown for significant SNO changes among all TCA enzymes with Cys nitrosylation sites, indicated as “modification sites,” as determined by MS. Green values indicate that inhibition in metabolic flux, as determined in isotopic labeling experiments (see Figure 2) was reversed by the NOS inhibitor L-NAME, consistent with the notion that S-nitrosylation mediated this inhibition. (**B**) Schema showing effects of S-nitrosylation (SNO) of TCA cycle enzymes in isogenic WT/Control and AD mutant hiN. AD-hiN display basal partial inhibition of the TCA cycle at the Aco/IDH steps. This is consistent with data that at least one of these enzymes is S-nitrosylated in AD-hiN (see panel E, below). S-Nitrosylation of αKGDH results in more major enzyme inhibition, as evidenced by reversal by L-NAME, and hence significant kinetic inhibition of flux through the TCA cycle at that point in the cycle. The addition of L-NAME increased the kinetic rate 1.4-fold at the aKGDH/SCS step in the AD-hiN, back to that of WT/Control-hiN, indicating that S-nitrosylation had slowed the rate by inhibiting enzyme activity. Note that the action of αKGDH also supplies NADH to the ETC for production of ATP. We have previously performed activity assays showing that S-nitrosylation of Aco, IDH, αKGDH and its DLD subunit can inhibit their activity (Doulias et al., 2021). Additionally, inhibition was observed after SNO at the level of PDH, which is S-nitrosylated via its DLD subunit in AD-hiN and human AD brain. Aco, aconitase or aconitate hydratase; CS, citrate synthase; DLD, dihydrolipoyl dehydrogenase (subunit 3 of αKGDH and PDH); ETC, electron transport chain; FH, fumarase or fumarate hydratase, IDH, isocitrate dehydrogenase; αKGDH, α-ketoglutarate dehydrogenase; MDH, malate dehydrogenase (mitochondrial) ME2, malic enzyme 2 (cytoplasmic); PC, pyruvate carboxylase; PDH, pyruvate dehydrogenase; SDH, succinate dehydrogenase (also complex II of the ETC); VLCAD, very long-chain acyl-CoA dehydrogenase. (**C**) Biotin-switch assay to confirm S-nitrosylation of TCA cycle enzymes in AD-hiN vs. isogenic WT/Control hiN. hiN cell lysates were subjected to biotin-switch assay for detection of SNO-αKGDH subunit 1 and standard immunoblot for total input αKGDH subunit 1. (**D**) hiN cell lysates were subjected to biotin-switch assay for detection of SNO-DLD (subunit E3 of αKGDH) and standard immunoblot for total input DLD. (**E**) hiN cell lysates were subjected to biotin-switch assay for detection of SNO-IDH α-subunit (SNO-IDH3Aα) and standard immunoblot for total IDH3Aα. For each biotin-switch assay, a representative blot is shown at left and a histogram of the densitometry of the bands at right. The addition of L-NAME to inhibit NOS or the omission of ascorbate serve as negative controls for the biotin-switch assay. Data are mean + SEM, n = 3, *p < 0.05; **p < 0.01 by ANOVA with Tukey’s correction.

To confirm the finding of S-nitrosylated TCA enzymes in human AD brains, we employed a second MS platform in the current study using a chemoselective-organomercury column to enrich for S-nitrosylated proteins followed by localization of the SNO-sites with MS (Doulias et al., 2010; Doulias et al., 2013b). The analysis on 3 male AD brains vs. 3 control male brains (**Table S1**, EXCEL spreadsheet 1, labeled “Human Brains Information”) showed, as expected, that many proteins were S-nitrosylated differentially in AD brains than in controls (all detected SNO-peptides and sites shown in **Table S1**, EXCEL spreadsheets 2-13 with Venn diagram summary in spreadsheet 9). Interestingly, gene ontology (GO) analysis of biological processes affected by SNO-proteins showed that TCA cycle enzymes and related metabolic pathways were again among the top hits (**Table S1**, EXCEL spreadsheet 13, labeled “GO_BP_AD,” entries highlighted in yellow). In addition to detection of SNO-Aco, the chemoselective-organomercury/MS dataset revealed that the only S-nitrosylated TCA enzymes that were found exclusively in AD brains and not in Controls were isocitrate dehydrogenase (IDH α-subunit) and αKGDH subunit 1 (**Table S1**, EXCEL spreadsheet 5, labeled “Unique to AD,” highlighted in yellow). Notably, in the initial *SNO*TRAP/MS dataset, although IDH α-subunit was also found to be S-nitrosylated in more AD brains than in non-AD controls, the mean ratio of spectral counts did not reach significance due to the variance between samples (ratio AD/Control = 1.67, p = 0.113 by ANOVA) (**Figure 1A**). Using the chemoselective-organomercury/MS approach, SNO-DLD could not be detected under our conditions so this enzyme could not be compared between the two datasets. Overall, the two detection methods for SNO-proteins were in agreement for identifying increased S-nitrosylation of TCA cycle enzymes although some details of the SNO-sites differed between the two techniques, as might be expected for different methods that complement one another. In the case of the *SNO*TRAP/MS approach, spectral counts could be used for semiquantitative comparison between AD and Controls to identify SNO-enzymes in the TCA cycle predominantly found in AD (Aco and DLD, representing subunit E3 of αKGDH) (**Figure 1A**). For the chemoselective-organomercury/MS technique, spectral counts for semiquantitative analysis are not available because we do not have multiple peptides for all the proteins, but some TCA SNO-enzymes (e.g., IDH α-subunit and αKGDH subunit 1) were found exclusively in AD human brains and not in Control brains (**Table S1**, EXCEL spreadsheet 5, labeled “Unique to AD”).

Next, to model the effect of these SNO-TCA cycle enzymes in a tractable system for mechanistic studies, we compared presenilin 1 (PSEN1) mutant AD-hiN to isogenic WT/Controls. Previously, PSEN1 mutant AD-hiN have been shown to manifest several features of human AD brains, including an increased β-amyloid (Aβ)42/40 ratio (Paquet et al., 2016; Ghatak et al., 2019; Ghatak et al., 2021). Here, also similar to results in AD human brains, by biotin-switch assay we found the largest increases in AD-hiN compared to isogenic WT/Controls occurred in SNO-αKGDH (subunit 1) and SNO-DLD (E3 subunit of αKGDH). (**Figures 1C** and **D**). Additionally, the IDH α-subunit was found to be significantly S-nitrosylated in AD-hiN over isogenic WT/Controls (**Figure 1E**). Notably, we have previously demonstrated that formation of SNO-TCA enzyme at levels similar to those found here (as judged by the ratio of S-nitrosylated enzyme to total enzyme) significantly inhibits enzyme activity (Doulias et al., 2021; Doulias et al., 2023). Taken together, these findings suggest that under similar conditions, metabolic flux should be inhibited for these TCA cycle enzymes. Hence, we next performed metabolic flux experiments.

### Metabolic flux assessment of defects in TCA cycle function in AD-hiN

Basal neuronal energy is thought to be generated by the TCA cycle after influx of lactate into neurons that is supplied by astrocytes (Magistretti and Allaman, 2015, 2018; Pellerin and Magistretti, 1994). However, during periods of intense stimulation, neurons can use glucose directly (Diaz-Garcia et al., 2017). Nonetheless, the lactate shuttle from astrocytes to neurons is critical for energy production to support normal synaptic function (Li and Sheng, 2022; Cheng et al., 2022). Importantly, recent evidence has shown that for prolonged cognitive loads, the use of lactate may be indispensable compared to glucose, suggesting that prevention of cognitive decline as seen in dementia, is dependent on the TCA cycle (Dembitskaya et al., 2022). To test this in AD-hiN vs. WT/isogenic-hiN, we supplied isotopically-labeled C^13^-lactate as the sole source of energy to hiN cultures for metabolic analysis. Our use of isogenic controls minimized the potential obfuscating effect of genetic background (Ghatak et al., 2019; Ghatak et al., 2021; Doulias et al., 2023).

For metabolic flux experiments, hiN were grown in cultures lacking astrocytes to which labeled lactate was added in order to simulate lactate supplied by astrocytes to neurons. Lactate is subsequently taken up by neurons and converted to pyruvate to supply the TCA cycle, whose individual enzyme activity and kinetic rate constants (i.e., flux through the system) can be monitored via monitoring the labeled metabolites (Ruiz et al. 2015; Cordes and Metallo, 2019; Torrini et al., 2022), as demonstrated here (**Figure 2**) and in Doulias et al. (2023) in Figure S1. As we and others have previously described, three critical principles are employed in interpreting these metabolic flux data (Fernandez-Garcia et al., 2020; Doulias et al., 2023):

1. Inhibition of a specific enzyme results in an increase in ^13^C-labeled substrate and a decrease in ^13^C-labeled product, resulting in an increased ratio of substrate/product, as measured in lysates by liquid chromatography-mass spectrometry (LC-MS) of TCA metabolites.
2. Relief of enzyme inhibition results in a decrease in the ratio of substrate to product back towards baseline. Here, to test for the effect of reversal of protein S-nitrosylation on relieving blocks in enzymatic activity, we used the nitic oxide (NO) synthase inhibitor, L-N^G^-nitro arginine methyl ester (L-NAME). L-NAME is known to prevent generation of reactive nitrogen species (RNS) related to NO that are known to be dramatically increased in AD brain and hiN in response to oligomeric Aβ (Talantova et al., 2013) and mediate aberrant protein S-nitrosylation of critical cysteine residues in or near active sites of multiple enzymes that affect metabolic flux through the TCA cycle (Doulias et al., 2021; Nakamura and Lipton, 2017; Doulias et al., 2023). Additionally, the converse is also true – exposure to an NO donor/transnitrosylating agent such as S-nitrosocysteine (SNOC) should increase protein S-nitrosylation of TCA enzymes (Doulias et al., 2021; Nakamura and Lipton, 2017).
3. For C^13^-lactate supplying the TCA cycle, the first two ^13^C-labeled carbons in the TCA enzyme substrates and products (labeled M+2) are likely supplied to the TCA cycle directly by lactate via one turn through the cycle. In contrast, if additional carbons are considered they may come from ancillary pathway metabolism (as shown in Figure S1 of Doulias et al., 2023) (Buescher et al., 2015; Butterfield and Halliwell, 2019; Jang et al., 2018; Lewis et al., 2014; Ruiz et al., 2015; Cordes and Metallo, 2019; Torrini et al., 2022). Therefore, data shown here represent raw data, after correction for background atmospheric C^13^ and normalization for mass, labeled as M+2 ME (molar enrichment) for each substrate and product (**Figure 2**, *left-hand panels*). The ratio of substrate/product is also shown, labeled M+2 MER (**Figure 2**, *right-hand panels*). Total ME and MER data for each pair of metabolites is shown in **Figures S1-S4**.

**Figure 2.**
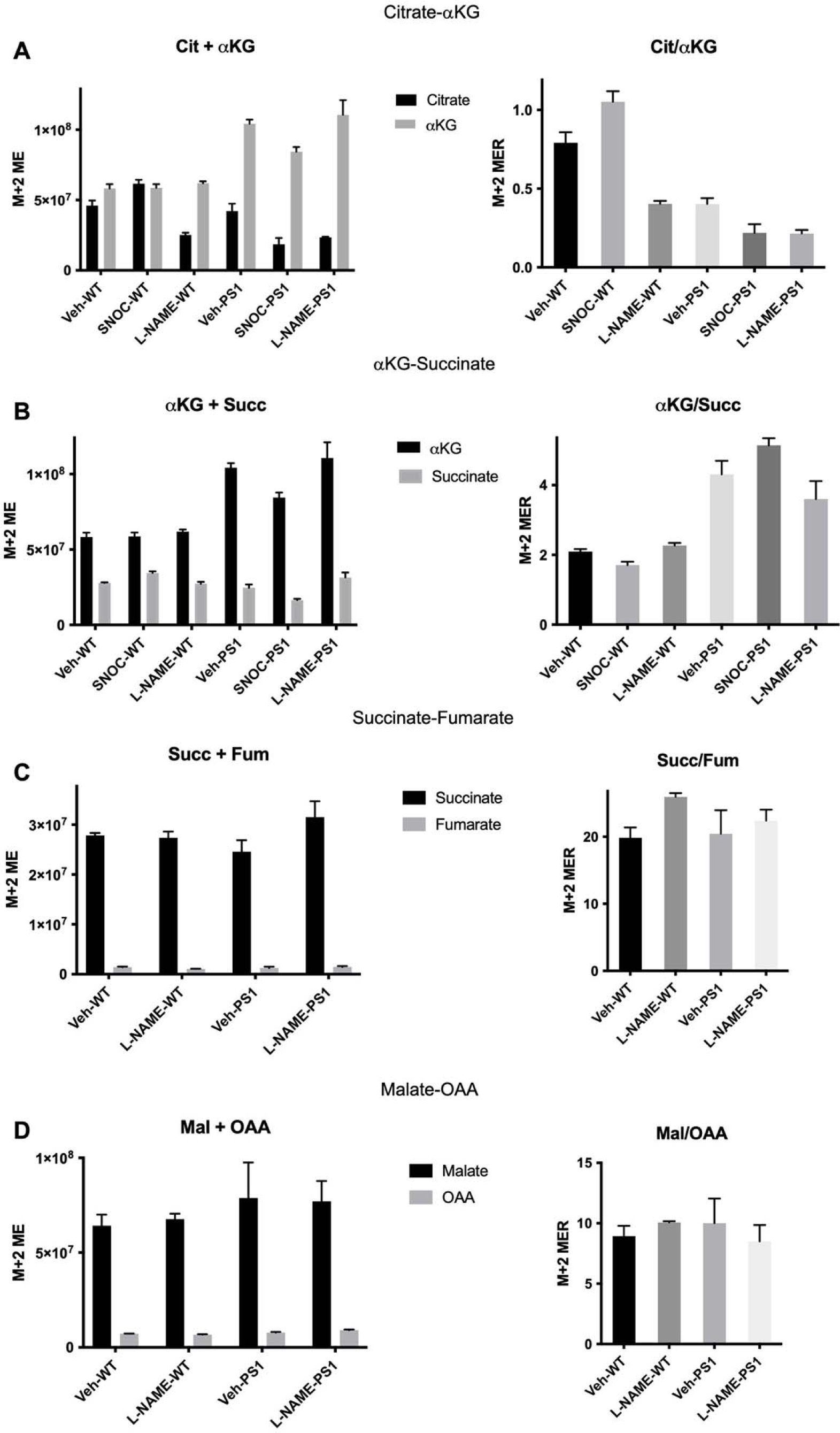
Metabolic flux analysis of TCA cycle enzymes. AD hiN (vs. isogenic WT/Control hiN incubated with ^13^C lactate. *Left-hand panels* show molar equivalents of M+2 isotopologues (M+2 ME) of individual metabolites; *right-hand panels* show corresponding molar equivalent ratios (M+2 MER) of substrate-product couples. (**A**) Substrate and product, citrate (Cit) and α-ketoglutarate (αKG), respectively, of reactions carried out by the aconitase/isocitrate dehydrogenase (Aco/IDH) segment of the TCA cycle. (**B**) α-KG and succinate (Succ) are substrate and product, respectively, of reactions carried out by α-ketoglutarate dehydrogenase/succinyl coenzyme-A synthetase (αKGDH/SCS) segment of the TCA cycle. (**C**) Succ and fumarate (Fum) are substrate and product, respectively, of succinate dehydrogenase (SDH). (**D**) Malate (Mal) and oxaloacetate (OAA) are substrate and product, respectively, of mitochondrial malate dehydrogenase (MDH). Vehicle (Veh); S-Nitrosocysteine (SNOC), 100 µM; L-N^G^-Nitro arginine methyl ester (L-NAME), 1 mM. See also **Figures S1-S4**.

### Inhibition of metabolic flux at the level of aconitase and isocitrate dehydrogenase

Looking at the citrate (Cit) to α-ketoglutarate (αKG) ratios (**Figure 2A**, *right-hand panel*), the metabolic flux experiments show the ratio decreases in WT/Control-hiN after exposure to L-NAME, consistent with basal inhibition by S-nitrosylation at the level of Aco/IDH that is relieved by NO synthase inhibition (**Figure 2A**, *right-hand panel*, p = 0.0001). In fact, in accord with these results, in both control human brains and in WT/Control-hiN, some level of basal S-nitrosylation was observed for Aco and/or IDH (**Figures 1A** and **1E**, and **Table S1**, EXCEL spreadsheet 6, labeled “Shared Proteins and Sites,” lines 15-17, and 164-168). As expected, exposure to SNOC, which would further increase the level of S-nitrosylation, further enhanced the Cit/αKG ratio (**Figure 2B**, *right-hand panel*, p = 0.0025).

Interesting, for AD-hiN compared to WT/Controls, the Cit/αKG ratio was decreased (**Figure 1B**, *right-hand panel*, p = 0.0001). Inspection of the individual Cit and /αKG levels revealed the explanation for this (**Figure 1B**, *left-hand panel*), as the difference was mainly in the increase in αKG level (p < 0.0001), with virtually no change in the Cit level in AD-hiN compared to WT/Controls. This finding is best explained by the presence of a more prominent block downstream in the TCA cycle in the AD-hiN, as discussed below. In this case, L-NAME would further decrease the Cit/αKG ratio in AD-hiN, (**Figure 1B**, *right-hand panel*, p = 0.0189), as the smaller block at the level of Aco/IDH would be relieved. The fact that SNOC also decreased the Cit/αKG ratio somewhat in AD-hiN(**Figure 1B**, *right-hand panel*, p = 0.0205) may indicate that the high levels of NO present at baseline in AD-hiN coupled with the addition of the NO donor may create such high levels of S-nitrosylating species that another level of block upstream from Aco/IDH develops that limits citrate production, e.g., via inhibition of pyruvate dehydrogenase (PDH, **Figure 1B**), which contains a DLD subunit that is S-nitrosylated to a significant degree (p = 0.026) in human AD brain over control (**Figure 1A**).

### Inhibition of metabolic flux at the level of **α**-ketoglutarate dehydrogenase/succinyl coenzyme-A synthetase

At the next steps in the TCA cycle, i.e., at the level of α-ketoglutarate dehydrogenase/succinyl coenzyme-A synthetase (αKGDH/SCS), as reflected in the αKG/succinate (Succ) ratios (**Figures 1B** and **2B**), the effect of AD-hiN compared to WT/Control was most prominent in inhibiting flux. Increases in Succ and αKG, with an overall increase in the αKG/Succ ratio at this stage in AD-hiN (**Figure 2B**, *right-hand panel*, p < 0.0001) and to an even greater degree after SNOC (p < 0.0001), may reflect a relative block at the level of αKGDH/SCS induced by NO-related species resulting in protein S-nitrosylation and resulting inhibition of enzyme activity. Consistent with this notion, after the addition of L-NAME, the ratio of a αKG/Succ was not significantly affected in WT/Control hiN but decreased in AD-hiN such that it was no longer significantly different from WT/Control + L-NAME (**Figure 2B**, *right-hand panel*). The latter reflects partial relief by L-NAME of inhibition from S-nitrosylation at the level of αKGDH since AD-hiN were shown to manifest increased S-nitrosylation of αKGDH subunit 1 and its E3 subunit DLD (**Figures 1C** and **D**), similar to that found in AD brains (**Figure 1A** and **Table S1**, EXCEL spreadsheet 5, labeled “Unique to AD”).

### Lack of inhibition of metabolic flux at the level of succinate dehydrogenase

Reflecting flux at the level of succinate dehydrogenase (SDH), AD-hiN and WT/Control hiN displayed similar succinate/fumarate (Succ/Fum) ratios (**Figure 2C**, *right-hand panel*). The minor effect of L-NAME apparently increasing this ratio in WT/Control hiN did not reach statistical significance. The fact that we found some evidence for S-nitrosylation of mitochondrial SDH in both control and AD human brains (e.g., **Table S1**, EXCEL spreadsheet 6, labeled “Shared Proteins and Sites,” lines 265, 266) suggests that some amount of regulation by S-nitrosylation might occur at this level, but the finding that L-NAME had little or no influence on the Succ/Fum ratio argues that the effect is a relatively minor one.

### Lack of inhibition of metabolic flux at the level of malate dehydrogenase

Similarly, distally in the TCA cycle, at the level of MDH, we observed no significant changes in the ratio of malate to oxaloacetate (Mal/OAA) when comparing AD-hiN to WT/Control hiN or when adding L-NAME to prevent S-nitrosylation. Interestingly, we did find evidence for S-nitrosylation at cysteine 93 of mitochondrial MDH in control and AD human brains (**Figure 1A** and **Table S1**, EXCEL spreadsheet 6, labeled “Shared Proteins and Sites,” line 184); note, however, that in AD human brains, S-nitrosylation at cysteine 93 decreased and a new site at cysteine 212 was detected (**Table S1**, EXCEL spreadsheet 8, labeled “Shared Proteins New Sites,” line 20; spreadsheet 10, labeled ‘”Shared Sites p < 0.05,” line 10; spreadsheet 11, labeled “TCA Enzymes,” line 17). Nonetheless, the fact that L-NAME displayed little effect on the Mal/OAA ratio argues that the effect of S-nitrosylation at these sites on MDH does not significantly affect enzymatic activity in our cell-based model. This result emphasizes why S-nitrosylation sites must be evaluated in the context of functional effects on activity for each enzyme and for an effect on flux through the TCA cycle at that step.

In summary, in AD-hiN inhibition of metabolic flux at the level of αKDGH/SCS was most prominent compared to WT/Control hiN and at least partially reversed by L-NAME. These findings are consistent with the notion that this is the most prominent site of inhibition in the TCA cycle by S-nitrosylation in AD-hiN. A minor inhibitory effect was observed at the level of Aco/IDH in both AD-hiN WT/Control hiN, indicating that it probably represents a normal regulatory mechanism at this stage of the TCA cycle. Kinetic modeling of the rate constants of the various TCA cycle enzymes (see METHOD DETAILS) confirmed that inhibition of the TCA cycle at αKDGH/SCS was the most prominent step affected by aberrant S-nitrosylation in AD-hiN compared to WT/Control-hiN. Notably, the rate constant for αKDGH/SCS increased significantly for AD-hiN values after addition of L-NAME (back to 100% from ∼70% in the AD-hiN and ∼60% in SNOC-exposed AD-hiN relative to the WT/Control-hiN value), signifying relief of inhibition and consistent with the notion that this was the major site of blockade in the TCA cycle by S-nitrosylation (**Figure 1B**).

### Mitochondrial bioenergetics in AD-hiN vs. WT/Control-hiN assessed by Seahorse metabolic flux analysis

For an enzymatic deficiency such as that induced by aberrant protein S-nitrosylation to manifest at a physiological level, it must be severe enough to restrict metabolic flux through an entire critically important pathway. In other words, it has to either affect the rate-limiting step or cause a shift of rate-limiting step to the enzyme in question. In the case of TCA cycle enzymes, the primary pathological scenario would be a decrease in the energy-transducing capacity of mitochondria that negatively affects their ability to produce ATP.

To assess the bioenergetic competence of mitochondria in hiN, we employed the so-called mitochondrial stress test assay using the Seahorse XF^e^96 metabolic flux analyzer. Interpretation of mitochondrial stress test data and optimized test format are discussed elsewhere (Ryan et al., 2013; Andreyev et al., 2015). Briefly, deficits in mitochondrial respiration are critically associated with neuronal dysfunction in AD. Therefore, we evaluated mitochondrial function in both AD-hiN and WT/Control-hiN. Using the Seahorse platform to measure OCR, we found a significant deficit in maximal rate of mitochondrial respiration in AD-hiN compared to WT/Control, as assessed after repeated applications of 2,4-dinitrophenol (DNP), which uncouples OXPHOS from the electron transport chain (ETC) (**Figures 3A** and **3B**). AD-hiN manifested a deficiency in spare respiratory capacity of ∼50% compared to WT/Control-hiN, even when corrected for cell number in the assay (**Figures 3A-3C**). Both genotypes, however, still possessed some degree of spare respiratory capacity (excess OCR in maximally DNP-stimulated state compared to the basal rate at the start of the experiment in **Figure 3A**). This suggests that the resting/unchallenged AD-hiN are not energy-deficient but cannot match the response of WT-Corrected hiN to an increased energy demand, in this case mimicked by the uncoupler DNP. In a physiological setting, the ramped-up energy demand may arise from increased firing rate or synaptic activity.

**Figure 3.**
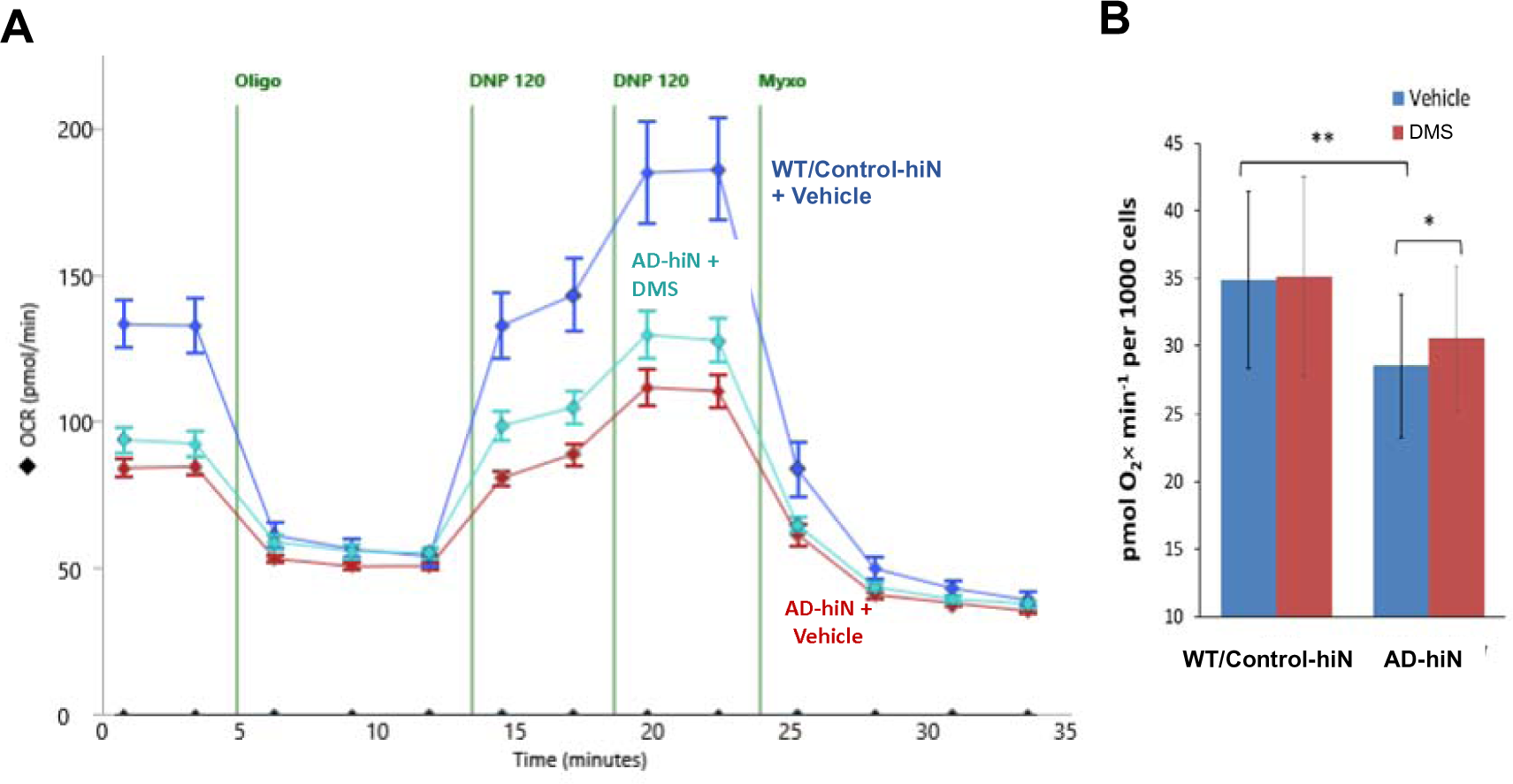
Respiratory defect in AD-hiN compared to WT/Control-hiN and partial rescue with dimethyl succinate. (**A**) Representative experimental run in Seahorse Flux Analyzer with 8-week terminally differentiated AD-hiN and WT/Control-hiN (10 wells per experimental group). OCR, oxygen consumption rate. Injections shown by vertical lines: Oligo, 2 μg/ml oligomycin; DNP, 120 μM 2,4-dinitrophenol; Myxo, 2 μM myxothiazol; DMS, 5 mM dimethyl succinate added 20 min prior to the run. (**B**) Respiratory capacity representing maximal uncoupler-induced OCR per 1,000 cells attained after 4 sequential additions of the uncoupler DNP. Data are mean ± SEM determined in replicate cultures of pure neurons (n = 10 wells per group in a single plate of hiN per experiment, with data from 14 separate experiments obtained in separate hiPSC differentiations. *p < 0.05, **p < 0.01 by two-tailed paired Student’s t-test.

In the resting as well as the stressed state in AD-hiN, part of the bioenergetic burden is shifted from functionally challenged OXPHOS to compensatory glycolysis (a Warburg-like effect), as observed also in AD neurons that have been directly converted from AD patient fibroblasts (Traxler et al., 2022). As an example of this, the ECAR data from the mitochondrial stress test show increased acidification rates in WT/isogenic-hiN over AD-hiN, indicative of increased glycolysis in the WT/isogenic-hiN over the AD-hiN; in the initial basal state ECAR, however, was higher in AD-hiN over WT/isogenic-hiN, indicating some degree of compensation in the AD-hiN (**Figure S5A**). Assessment of ATP production rates in the basal state, indicative of ATP demand by the cells, shows virtual independence of total ATP demand with respect to genotype (**Figure S5B**), with an almost twofold increase in reliance on aerobic glycolysis (**Figure S5C**). It should be noted, however, that the contribution of glycolysis compared to OXPHOS remains very minor (∼6% and ∼11% for WT/Control-hiN and AD-hiN, respectively).

Further comparing the capacity of glycolysis vs. OXPHOS in both the WT/Control-hiN and AD-hiN, we found that ATP production was primarily through OXPHOS (i.e., TCA cycle/ETC pathways) rather than glycolysis (**Figure S6**). Accordingly, there was a direct correspondence between the lower OCR values of AD-hiN compared to WT/ corrected-hiN (**Figure 3B**) and the significantly decreased (p < 0.01) ATP-producing capacity of OXPHOS in AD-hiN vs. WT/Control (**Figure S6**). Perhaps unexpectedly, the capacity of glycolysis to produce ATP in AD-hiN was slightly but significantly elevated (p < 0.001) compared to WT/Control-hiN, suggesting that, beyond the mass action-driven compensatory activation, there is an active upregulation of the pathway in AD-hiN.

### “Substrate bypass” in the TCA cycle results in improved bioenergetics in AD-hiN

Next, we reasoned that by supplying excess substrate to bypass the step in the TCA cycle that was most inhibited via aberrant S-nitrosylation in AD-hiN, namely, the production of succinate by αKDGH/SCS, we might be able to improve the overall bioenergetics in these neurons. We therefore tested the ability of a membrane-permeant form of succinate, dimethyl succinate (DMS) (Selak et al., 2005; Chouchani et al., 2014), to rescue the respiratory deficiency in AD-hiN. Importantly, supplying a pro-drug like DMS that is metabolized to succinate not only provides substrate to the next enzymatic step in the TCA cycle, SDH, but also to the ETC since SDH also functions as complex II in the ETC (**Figure 1B**). As predicted, we found using the Seahorse platform that acute application of DMS to AD-hiN produced partial recovery of respiratory capacity (**Figure 3A**). Over multiple experiments, respiratory capacity of the AD-hiN was ∼20% lower than WT/Control-hiN, and DMS treatment resulted in recovery of about a third of the deficit (**Figure 3B**). This result represents proof-of-principle that bypass of the block in the TCA cycle could rescue the bioenergetics of AD neurons.

As a further indication that DMS enhanced the bioenergetics of AD-hiN, we monitored the NADH/NAD^+^ ratio using a specific autofluorescence technique (see METHOD DETAILS) and found a relative increase in NADH in AD-hiN after DMS treatment (**Figure S7**). This finding is consistent with the notion that by bypassing the relative block at the αKGDH/SCS step where NADH is produced, additional NADH is generated more distally in the TCA cycle, e.g., at the malate to oxaloacetate step catalyzed by MDH (**Figure 1B**).

### “Substrate bypass” partially rescues synapses in AD-hiN

Loss of synaptic function is considered a primary determinant of AD pathology and cognitive decline (Terry et al., 1991; DeKoskey et al., 1990; Tzioras et al., 2023). Since the function and maintenance of synapses are energy-intensive processes important to cognitive activity and the mitochondrial TCA cycle in neurons is critically important in this regard (Dembitskaya et al., 2022; Tzioras et al., 2023), even the relatively modest bioenergetic defect observed under stress in AD-hiN may contribute to synaptic dysfunction and loss. Therefore, we monitored synapse number in cultured AD-hiN vs. WT/Control-hiN using immunocytochemistry for pre-and postsynaptic markers with high-content imaging on the automated ImageXpress Micro Confocal imaging platform (Molecular Dynamics, Inc.). We found a statistically significant deficit in synapse number in AD-hiN compared to WT/Control hiN (**Figures 4A** and 4B), reminiscent of the degree of synaptic loss reported in both transgenic AD mice and human AD brains (Terry et al., 1991; DeKoskey et al., 1990; Lipton et al., 2016; Tzioras et al., 2023). Notably, treatment of cultures for 48 hr with DMS offered statistically significant protection of synapse number in the AD-hiN (**Figure 4C**). DMS treatment increased synapse density in AD-hiN nearly 70% (calculated from 13% loss of synapses in AD-hiN compared to WT/Control hiN in **Figure 4B**, and 9% recovery of synapse number in AD-hiN with DMS in **Figure 4C**).

**Figure 4.**
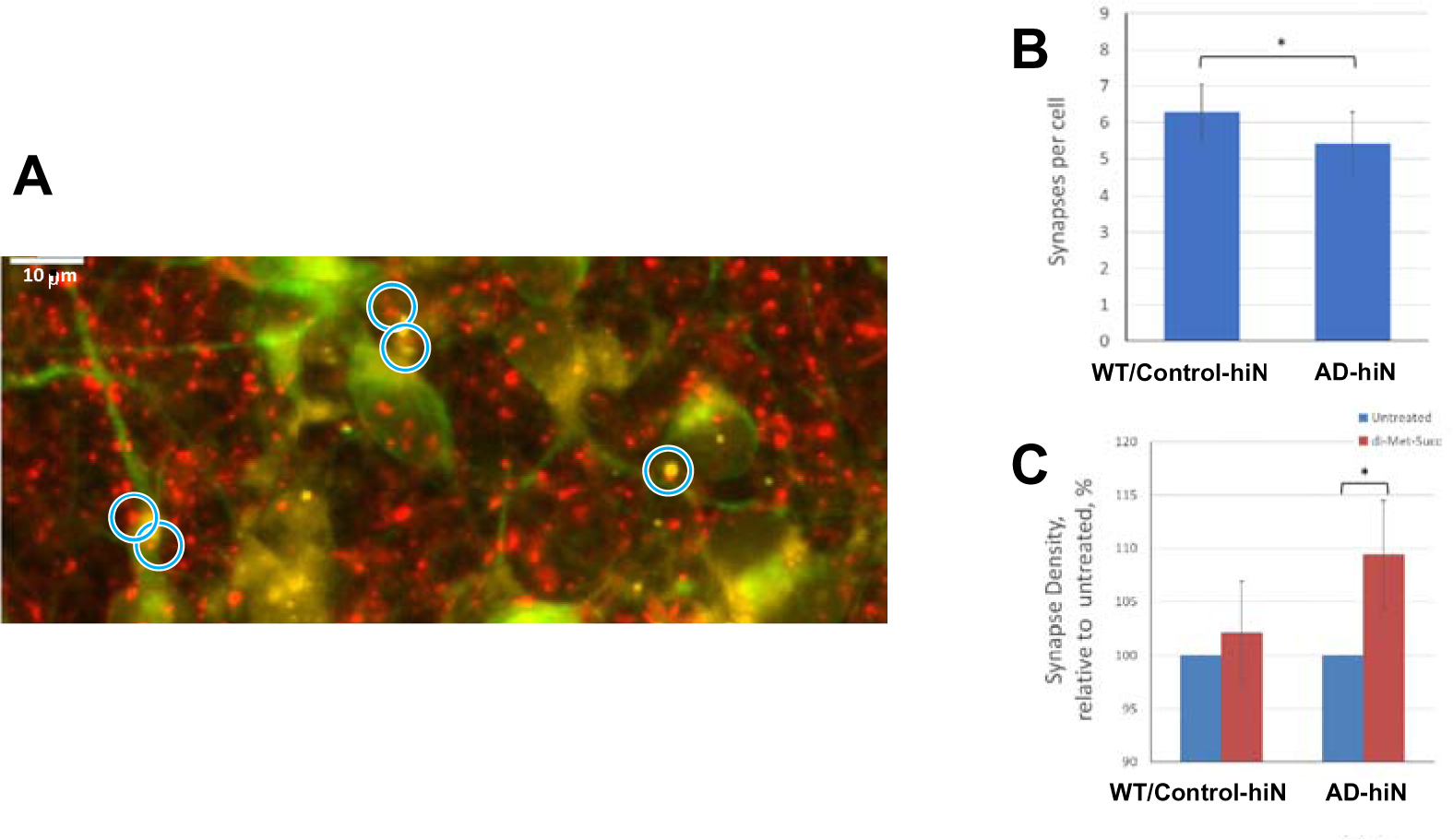
Quantification of synapse number in AD-hiN and WT/Control hiN. (**A**) Representative field of AD-hiN stained for the presynaptic marker Synapsin 1 (red), the postsynaptic marker Homer 1 (yellow), and the neuronal marker microtubule-associated protein 2 (MAP2, green). Synaptic punctae (blue circles) were identified by co-localization proximity of pre-and postsynaptic markers (see METHOD DETAILS). (**B**) Quantification of synaptic punctae per cell in a typical experiment demonstrating ∼14-15% loss of synapses in the AD-hiN compared to WT/Control-hiN. Values are mean ± SEM of 25 separate sets of hiN cultures, each represented by a 10-well group in a 96-well plate; *p < 0.05, by Wilcoxon signed-rank test. (**C**) Dimethyl succinate (DMS, 5 mM) treatment for 48 hr led to recovery of synaptic density relative to untreated in AD-hiN but had no significant effect on WT/Control-hiN. Data are mean ± SEM; *p < 0.05, by Wilcoxon signed-rank test (n = 25 plates assayed).

## DISCUSSION

While other publications have suggested a common feature in neurodegenerative disorders of aberrant S-nitrosylation of TCA cycle enzymes (Nakamura and Lipton, 2017; Doulias et al., 2021; Doulias et al., 2023), this is the first study to demonstrate that substrate-bypass of the major SNO-induced block in the TCA cycle can at least partially rescue energy supply and, critically, protect from synaptic damage. A recent review emphasized the importance of studying covalent modification that modulate the TCA cycle (Arnold and Finley, 2022). In this vein, we describe partial inhibition in flux through the TCA cycle in AD-hiN caused by aberrant protein S-nitrosylation prominently at the αKGDH step (subunit 3, DLD, in **Figure 1D**), which would normally produce NADH and convert αKG to succinate. NADH is thus supplied to complex I of the ETC, and succinate as substrate to SDH, which also functions as complex II of the ETC (**Figure 1B**). Therefore, inhibition of αKGDH results in both a decrease in substrate for SDH/complex II and also prevents NADH supply to complex I, thus limiting the production of ATP by mitochondria (Butterfield and Halliwell, 2019; González-Rodriguez et al., 2021; Jang et al., 2018; Lewis et al., 2014; Nakamura and Lipton, 2017). In this manner, formation of SNO-αKGDH would contribute to the bioenergetic compromise observed in AD-hiN by Seahorse metabolic flux analysis. Collectively, in both animal models and in hiPSC-based models, we and others have causally linked such bioenergetic compromise in mitochondria to the pathogenesis of AD and other neurodegenerative disorders by contributing to synaptic loss, and eventually to neuronal cell death (Cho et al., 2010; Cooper et al., 2012; Czaniecki et al., 2019; Gibson et al., 2010; Ryan et al., 2013; Ryan et al., 2018; Nakamura et al., 2021). The present study links these findings of compromised mitochondrial bioenergetics to TCA cycle compromise, precipitated by aberrant protein S-nitrosylation reactions observed in human AD brains.

Moreover, our concordant data between chemical S-nitrosylation in human AD brains and enzymatic inhibition in the TCA cycle in AD-hiN □ for each of the S-nitrosylation events in TCA cycle enzymes □ are consistent with the notion that this posttranslational modification is in large part responsible for the observed inhibition at the αKGDH step of the TCA cycle in human AD brains (Gibson et al., 2010). Furthermore, the fact that the nitric oxide synthase (NOS) inhibitor L-NAME could reverse the inhibition in metabolic flux monitored with isotope label at the αKGDH step in our AD-hiN cultures is consistent with the notion that NO-mediated S-nitrosylation is responsible for the abnormal inhibitory effect at this step in the TCA cycle that is observed in human AD brains. Of note, other recent findings in another model system using direct conversion of AD patient fibroblasts to neurons have also indicated a Warburg-like effect with TCA cycle shutdown in AD neurons (Traxler et al., 2022).

In terms of potential therapy, when we delivered a membrane-permeable form of succinate to bypass the relative block at the αKGDH enzymatic step to AD-hiN, we could partially rescue bioenergetic compromise as assessed by in the Seahorse metabolic flux analyzer and, most importantly, fully restore synapse number. The addition of succinate acts in a dual fashion by increasing the total pool of TCA cycle intermediates (anaplerotic action) and by bypassing compromised complex I of the ETC (as has been reported in AD brains (Eckert et al., 2011)). Maintenance of even the minimal segment of the TCA cycle metabolizing succinate to oxaloacetate that is not dependent on the activity of DLD-containing enzymatic complexes (PDH and α-KGDH), generates one NADH and one FADH_2_, which account for about one third of the stoichiometry of the intact TCA cycle. Interestingly, the degree of synaptic loss we observed in our AD-hiN cultures is similar to the level of loss previously reported in AD transgenic animal models and human AD patients (Terry et al., 1991; DeKoskey et al., 1990; Lipton et al., 2016; Tzioras et al., 2023), and we could virtually completely rescue this degree of synaptic loss, at least in our AD-hiN models. This not only serves as proof-of-principle that metabolic rescue can affect synaptic number but also suggests future therapeutic avenues that can be pursued for AD, especially in light of the fact that synaptic loss is arguably the best single correlate to cognitive decline in human AD patients (Terry et al., 1991; DeKoskey et al., 1990; Tzioras et al., 2023).

### Limitations of the study

This study is based on findings in postmortem human brain from AD patients and controls. While postmortem times were short (generally a few hours), there is always the possibility that agonal changes could influence the findings. That said, the fact that hiPSC-based cultures of neurons with AD mutation yielded similar results gives us added confidence in the findings. Moreover, the stress present in these culture systems may cause the neurons to age more quickly (Bhaduri et al., 2020), and thus provide a reasonable model of neurodegenerative diseases of aging like AD. Nonetheless, using cultured neurons, including AD-hiN, represent *in vitro* conditions, and thus inform on what is plausible rather than faithfully replicating *in vivo* conditions. In the present study, the use of pure hiN in the absence of astrocytes could be viewed as an artificial system. This approach, however, allowed the assessment of human cells from a patient with the disease process and comparison to isogenic, gene-corrected controls. The fact that several features of AD are faithfully reproduced in these AD-hiN cultures such as a hyperelectrical phenotype and synaptic damage (Ghatak et al., 2019; Ghatak et al., 2021) gave us increased confidence that aspects of AD such as synaptic loss could be reproduced and studied. Moreover, our finding of similar aberrant S-nitrosylation reactions in the TCA cycle in fresh postmortem human brain specimens from AD patients and AD-hiN supports the pathophysiological significance of our results. Thus, the hiPSC platform was useful for modeling metabolic and synaptic features of AD. Another critical consideration in the analysis of metabolic experiments is concern over which compartments are being analyzed. In our case, we were able to discern the location of enzymes as mitochondrial vs. cytoplasmic via their exact amino-acid sequences obtained from the MS data for the protein S-nitrosylation experiments since the mitochondrial and cytoplasmic versions of these enzymes are in general not identical. Moreover, for the metabolic flux experiments, by focusing on the M+2 analysis after heavy isotope labeling of lactate, we limited our conclusions primarily to data obtained after one turn through the TCA cycle in order to focus on events primarily driven by entry into the TCA cycle via lactate and to avoid ancillary pathways of entry.

## STAR★METHODS

### EXPERIMENTAL MODEL AND SUBJECT DETAILS

#### hiPSC cultures

Pluripotent cells (hiPSCs) were cultured and maintained in our laboratory using a protocol described previously (Ghatak et al., 2019; Ghatak et al., 2021). M146V/WT hiPSC lines bearing the PSEN1 (PS1) M146V mutation and the associated isogenic WT control (from the laboratory of Marc Tessier-Lavigne, Rockefeller University/Stanford University, and the NY Stem Cell Institute, NYC). Details regarding these lines have been previously published (Paquet et al., 2016) and they were re-karyotyped to ensure genomic stability. hiPSCs were plated on γ-irradiated human foreskin fibroblasts and cultured using medium containing 20% KSR and 8 ng/ml bFGF, changed daily, as we have described (Ghatak et al., 2019; Ghatak et al., 2021). The colonies were manually passaged weekly.

#### hiN neuronal differentiation

Differentiation of hiPSCs into cerebrocortical neurons (hiN) and detailed characterization of these cultures were performed, as we have described previously (Ghatak et al., 2019; Ghatak et al., 2021). Just before differentiation the colonies were dissociated into single cell suspension using accutase. To purify hiPSCs and remove fibroblast feeders, medium containing dissociated fibroblasts and hiPSCs was placed in gelatin-coated dishes. After adherence, supernatant containing purified hiPSCs was collected and re-plated at 4×10^4^ cells/cm^2^ on Matrigel (BD)-coated tissue culture dishes for differentiation. The resulting hiN were present in the absence of an astrocyte feeder layer; this was important so that the addition of isotopically labeled C^13^-lactate in the metabolic flux experiments mimicked the supply of lactate that would normally occur from astrocytes. For most experiments, 8-week terminally differentiated hiN were used.

#### Human AD and control brains

Two sets of human AD and control brains were used in this study. The detailed analysis of the first set, assessed for S-nitrosoproteins and SNO-sites with the S-nitrosothiol-specific triaryl phosphine probe, *SNO*TRAP, coupled to MS, was recently published (Yang et al., 2022). That study included autopsy-confirmed human brains with AD (n = 10 females and n = 10 males) and non-AD (*n* = 10 females and *n* = 10 males), matched for age, sex, education, and ethnicity (Yang et al., 2022). In a second set of human brains described here, we assessed S-nitrosoproteins and SNO-sites by MS after enrichment on a chemoselective-organomercury column (Doulias et al., 2010; Doulias et al., 2013b). This second analysis included autopsy-confirmed human brains with AD (n = 3 males) and controls (n = 3 males), matched for age, sex, education, and ethnicity (**Table S1**, EXCEL spreadsheet 1, labeled “Human Brains Information”). In all cases, the brain tissues were sampled from the prefrontal cortex (Brodmann area 10). All brain tissues were obtained from the University of California (UCSD) Medical Center and the San Diego VA Medical Center Brain Bank and were flash frozen at postmortem examination. The study was approved by the local Ethics Committee of both medical centers. Neurological diagnoses were performed independently by two experienced clinicians in alignment with the consensus criteria for AD and were confirmed neuropathologically by the presence of plaques and tangles in the face of neurodegeneration. All the tissues were stored at −80°C until use.

### METHOD DETAILS

#### Preparation of human brain tissues for mass spectrometry

Human brain tissues were homogenized on ice using a Teflon pestle and a Jumbo Stirrer (ThermoFisher) in freshly prepared lysis buffer (100 mM HEPES-NaOH, pH 7.7, 1 mM EDTA, 0.1 mM neocuproine, 1% Triton X-100, 20 mM IAM, 1% protease inhibitor cocktail, and 0.1% SDS). Homogenates were then centrifuged at 16,000 g for 15 min at 4 °C, and supernatants collected. Protein concentration was determined by the Bradford assay. One volume of 50 mM HEPES buffer (pH 7.7) was added to the supernatants, which were then centrifuged at 5,000 g for 25 min at 4°C using 10 K MWCO spin filters. The preparation was then stored at −80°C prior to use.

#### Assessment of S-nitrosylation sites in human AD and Control brains

For assessment of the S-nitrosoproteome in human brains using *SNO*TRAP labeling followed by MS, we used protocols that we recently published along with their results (Yang et al., 2022). In the present manuscript, the S-nitrosylated TCA enzymes in these brains are analyzed. In brief, *SNO*TRAP-labeling stock solutions prepared in acetonitrile (ACN) were added to samples at a final concentration of 1.5 mM to selectively convert SNO to stable disulfide-iminophosphorane. For negative controls, we doped the same volume of 40% ACN in 50 mM HEPES (pH 7.7). The samples were incubated at RT for 2 hr. After reaction, excessive reagents were removed with three washes of 50 mM HEPES, pH 7.7 buffer with 10 K MWCO filters, followed by trypsin digestion overnight at 37°C.

After enzymatic digestion into peptides, 200 μL Streptavidin agarose beads were added to each sample and incubated for 2 hr at RT with gentle shaking. To minimize non-specific binding, the beads were washed twice with 5-fold volumes of the following buffers in succession: Buffer I (100 mM NH_4_HCO_3_ (ABC), 150 mM NaCl, 1 mM EDTA, pH 7.4, containing 0.05% SDS, 0.1% Triton X-100); buffer II (100 mM ABC, 150 mM NaCl, 1 mM EDTA, pH 7.4 containing 0.1% SDS); buffer III (100 mM ABC + 0.05% SDS + 150 mM NaCl), buffer IV (100 mM ABC), and buffer V (50 mM HEPES, pH 7.7). Peptides bound to beads were then eluted with 10 mM TCEP (in 50 mM HEPES, pH 7.7) and alkylated with 100 mM NEM for 2 hr at RT. After alkylation, samples were desalted with Pierce C18 spin columns and stored at –80 °C prior to analysis. Sample preparation and storage were conducted in the dark.

The desalted peptides, dissolved in 0.1% FA, were then analyzed using a Q Exactive MS coupled to Easy-nLC 1000 (ThermoFisher Scientific, Waltham, MA) interfaced via a nanoSpray Flex ion source. Five technical runs were conducted for each biological replicate, i.e., from tissue of a single individual. Formic acid in water (0.1%) and in ACN (0.1%) were used as mobile phases A and B, respectively. Aliquots (3 μL) of the samples were injected onto a C18 pre-column (75 μm ID × 20 mm, 3 μm, Thermo Scientific) and separated by a C18 column (75 μm ID × 250 mm, 2 μm, Thermo Scientific) with a stepwise gradient (1% B for 10 □min, 1–60% B for 110 □min, and then 60-100% B for 10□ min) followed by a 10 min post-run at 1% B at a flow rate of 300□ nL/min. Mass spectra were acquired in data-dependent mode using the following settings: spray voltage, 2.2 kV; capillary temperature, 250 °C; S-lens RF level, 60%; no sheath and auxiliary gas flow; resolution 70,000; scan range 350–1800 Th. The 10 most abundant ions with multiple charge-states were selected for fragmentation with an isolation window of 2 Th, and a normalized collision energy of 28% at a resolution of 35,000. The maximum ion injection times for the full MS scan and the MS/MS scan were both 100 ms. The ion target values for the full scan and the MS/MS scans were set to 3□×□10^6^ and 1□×□10^5^, respectively. Xcalibur software was used for data acquisition.

For assessment of the S-nitrosoproteome in human brains using a second approach, organomercury-chemoselective enrichment of S-nitrosothiols, we followed a recently described protocol (Doulias et al., 2021). Experimental details for preparation and activation of columns, and reaction of homogenate with organomercury resin for S-nitrosocysteine enrichment have been presented previously (Doulias et al., 2010; Doulias et al., 2013b). For each group, five samples were analyzed with technical duplicates. For column washes, we used 50 bed volumes of 50 mM tris-HCl (pH 7.4), containing 300 mM NaCl and 0.5% SDS. Following this, 50 bed volumes of the same buffer containing 0.05% SDS were used. Next, columns were washed with 50 bed volumes of 50 mM tris-HCl containing 300 mM NaCl (pH 7.4), 1% Triton X-100, and 1 M urea. Columns were then washed with 50 bed volumes of the same buffer containing 0.1% Triton X-100 and 0.1 M urea, and finally 200 bed volumes of water. This was followed by on-column digestion into peptides; for this, columns were initially washed with 10 bed volumes of 0.1 M ammonium bicarbonate.

Bound proteins were then digested with Trypsin Gold (1 μg/mL) (Promega) that was added in a one bed volume of 0.1 M ammonium bicarbonate in the dark for 16 h at room temperature. Next, washing the resin with 40 bed volumes of 1 M ammonium bicarbonate (pH 7.4) containing 300 mM NaCl, and then 40 volumes of the same buffer without salt, 40 volumes of 0.1 M ammonium bicarbonate, and 200 volumes of deionized water. Bound peptides were eluted by treatment with one bed volume of performic acid in water (Doulias et al., 2010; Doulias et al., 2013a). To increase peptide recovery, the column was further washed with one volume of deionized water. The eluted peptides thus obtained were stored at −80 °C overnight, and then lyophilized and resuspended in a 300 μL volume of 0.1% formic acid. The peptides in the resuspended solution were put in low-retention tubes (Axygen) and the volume reduced to 30 μL by speed vacuum. Peptides were then desalted using Stage Tips (ThermoFisher) and transferred to a high-performance liquid chromatography (LC) vial for subsequent LC-tandem mass spectrometry (MS/MS) analysis of SNO-peptides using a Orbitrap Elite Hybrid Ion Trap-Orbitrap Mass Spectrometer (ThermoFisher). As previously described (Doulias et al., 2013a), we determined the S*-*nitrosoproteome by identifying S-nitrosylated peptides and proteins for each sample group.

Three different biological samples per group were each analyzed in duplicate. The last step for the organomercury-based site-specific identification of S-nitrosylated cysteine residues consists of using mild performic acid to release the bound peptides selectively and quantitatively. Importantly, the performic acid oxidizes the cysteine thiols to sulfonic acid, thereby generating a unique MS signature that permits site-specific identification of the modified cysteine residues as >96% of cysteine-containing peptides were detected with the sulfonic acid modification in the 20 samples analyzed. For the final reporting table all peptides detected in the duplicate samples for each condition were grouped (Raju et al., 2015). Note that the organomercury-chemoselective method identifies the sites of S-nitrosylation by detecting peptides with modified cysteine residues. Since, like other protein posttranslational modifications, S-nitrosylation is sub-stoichiometric, some peptides will not be captured and detected. In this regard, the presence of a particular peptide in one biological condition (e.g., Control brains) and its absence from another can be attributed to the following possible reasons: The cysteine residue is not S-nitrosylated, or its S-nitrosylation level is below the detection limit of the method.

#### MS data processing and statistical analysis of human brain S-nitrosoproteome

For assessment of the S-nitrosoproteome analyzed by the *SNO*TRAP/MS method, Agilent Spectrum Mill MS proteomics Workbench B.06 was used for peak list generation, database searching, label-free semiquantitative assessment, and false discovery rate (FDR) estimation. Parameters for data extractions were as follows: Precursor MH^+^ 300 – 8000 Da, scan time range 0-200 min, sequence tag length > 1, merge scans with same precursor *m/z* ±30s and ±0.05 *m/z*, default for precursor charge, true for find ^12^C precursor, MS noise threshold 100 counts. MS/MS spectra were searched against the human SwissProt protein database (downloaded on 06/18/2019) with ±10 ppm precursor ion tolerance and ± 20 ppm fragment ion tolerance. The search included variable modifications of methionine oxidation, protein N-terminal acetylation, deamidation of asparagine, and fixed modification of N-ethylmaleimide on Cys. For both peptide identification and protein polishing, the FDR was set to 1%. Peptide identifications were accepted only if the following confidence thresholds were met: Minimal peptide length was set to five amino acids, and a maximum of two missed cleavages was allowed. The MS/MS spectra were inspected manually to validate the peptide/protein identifications. In addition, for proteins detected in one group but not another, the raw LC/MS data files for the latter were searched manually for the presence of protein-related peptides detected in the former. The MS raw data have been deposited in the ProteomeXchange Consortium (http://proteomecentral.proteomexchange.org) via the PRIDE partner repository with the dataset identifier PXD042436.

For S-nitrosoproteins identified in brain with the *SNO*TRAP/MS technique, the precursor intensities of SNO-proteins, estimated in Spectrum Mill by combining precursor intensities of the constituent peptides in MS1 spectra, were used to calculate fold-change quantification of the common SNO-proteins detected in AD and Control groups, with fold-change defined as total-ion intensity AD sample divided by total-ion intensity of respective Control. Statistical evaluation of nanoLC-MS label-free differential data was performed after the application of data imputation to reduce the number of missing values. Missing values were replaced by this value according to the local Minimum method (van Ooijen et al., 2018). A fold change of >2 was considered increased expression, while fold change of <0.5, decreased expression; these cut-offs corresponded to a p-value < 0.05, as we have described previously (Seneviratne et al., 2016). For comparison of quantifications of SNO-proteins obtained with Spectrum Mill, an ANOVA followed by a post hoc Fisher’s PLSD test for multiple comparisons, with a difference of at least p < 0.05 considered statistically significant.

For S-nitrosoproteins enriched by the organomercury-chemoselective method, after column digestion to generate peptides and elution of peptides, the trioxidation of SNO-cysteine residues was used to identify the site of modification. Raw MS data were analyzed with MaxQuant open-source software using cysteine trioxidation and methionine dioxidation as differential modifications. We report the S-nitrosylation sites on the peptides/proteins based on the MaxQuant output with assessment of SNO-site presence or absence in specific proteins in AD vs. normal Control brain. In the case of the organomercury-chemoselective/MS method, we could not use spectral counting because we do not have multiple peptides for all the proteins. We therefore used the MS1 peak intensity of a peptide to compare between Control and AD brain. However, this comparison does not allow us to report on the most important aspect-occupancy of the sites. Occupancy is the fraction of SNO modified to unmodified peptide. Previously we found that the majority of SNO sites are not occupied by other cysteine modifications after organomercury-chemoselective enrichment (Doulias et al., 2010; Doulias et al., 2013b); thus, we consider occupancy only to unmodified peptide.

#### Bioinformatic analysis of S-nitrosoproteome data

To analyze functional enrichment of gene ontology (GO) processes and pathways for the human AD vs. Control brains characterized by *SNO*TRAP/MS, we uploaded the SNO-protein IDs into MetaCore^TM^ software (version 19.4 build 69900). Reactome pathway analysis was performed using STRING (version 11.0) (http://string-db.org) (Mi et al., 2013). High reliability interactions (score > 0.7) were analyzed. To visualize protein-protein interactions, we used Cytoscape plug-in software (version 3.7.2). FDR values <0.1 were accepted for the analyses using the aforementioned software platforms. The gene names for the unique proteins identified for each of the six conditions were submitted to Gene Ontology http://geneontology.org and analyzed for functional processes or cellular component (localization) enrichment.

Following S-nitrosoproteome identification in human brains by organomercury-chemoselective enrichment/MS, functional analysis was performed to characterize the biological pathways affected using gene ontology (GO) knowledgebase (2020-12-08 release) enrichment, pathway analysis using the Kyoto Encyclopedia of Genes and Genomes (KEGG release 96.0, October 1, 2020), and STRING protein-protein interaction network functional enrichment as we have described (Doulias et al., 2021).

#### Biotin-switch assay in hiN for S-nitrosylated TCA enzymes

As we have previously described, biotin-switch assays were performed with standard methods (Cho et al., 2009; Nakamura et al., 2021; Doulias et al., 2023). hiN lysates were prepared in HEN buffer [100 mM HEPES-NaOH (pH 7.4), 1 mM EDTA, 0.1 mM neocuproine] containing 150 mM NaCl, 1% NP-40, 0.5% deoxycholate, and 0.1% SDS. Next, free thiol groups were blocked with 10 mM S-methyl methanethiosulfonate (MMTS, Millipore Sigma, 208795) in HEN buffer at 42 °C for 20 min. Following the removal of excess MMTS via acetone precipitation, S-nitrosothiols were specifically reduced to free thiols using freshly prepared sodium ascorbate (10 mM, Millipore Sigma, 11140). Next, the newly formed free thiols were labeled with 1 mM Biotin-HPDP(WS) (Dojindo Molecular Technologies, SB17) at RT for 1 hr. The resulting biotinylated proteins were then enriched by binding to High Capacity NeutrAvidin Agarose beads (ThermoFisher Scientific, 29204). Purified biotinylated proteins were then eluted from the beads for immunoblot analysis.

For immunoblots, samples were subjected to Bolt 4-12% Bis-Tris mini protein gel electrophoresis (ThermoFisher Scientific, NW04122BOX) and transferred to Immobilon-FL PVDF membranes (Millipore Sigma, IPFL00010). Subsequently, the membranes were blocked using Intercept TBS blocking buffer (Li-Cor, 927-60001) and then incubated at 4 °C overnight with primary antibodies against αKGDH subunit 1 (Abcam, ab137773), lipoamide dehydrogenase (DLD, Abcam, ab133551), or IDH α-subunit (ProteinTech, 501730566). Following 3 washes with 1X TBS-T (Cell Signaling Technology, 9997S), the membranes were incubated with IR-dye-conjugated secondary antibody (IR-dye 800CW-conjugated goat anti-rabbit [1:15,000; Li-Cor, 926-32211]) at RT for 1 hr. Membranes were scanned with an Odyssey CLx infrared imaging system (Li-Cor). Image Studio software (Li-Cor) was used for densitometric analysis of immunoblots. Each set of biotin-switch assays and immunoblots was run on at least three independent biological samples.

#### Metabolic flux and related experiments

AD-hiPSCs and isogenic WT-hiPSCs were prepared and differentiated into hiN as described above. Two-thirds of the volume of the culture medium was exchanged biweekly with DAN medium, continuing for ∼5 weeks until experimentation. The last medium change contained 1 mM L-NAME for some wells for 24-hour pretreatment. All wells were then switched for 6 h to Deprivation Medium (± L-NAME) containing B27/N2 supplements, but without glucose/pyruvate/lactate. To ascertain how lactate is handled by the TCA cycle in AD-hiN vs. WT/Control-hiN, 1 ml of 3X (30 mM) concentrations of either ^12^C or ^13^C-lactate in Deprivation Medium was added to the appropriate wells. After 6 h labeling, conditioned medium was harvested and snap frozen. Cells were washed once with PBS, solution aspirated, and cells snap frozen on the plates. Plates and media were then harvested for analysis of labeled metabolites to assess metabolic flux.

Metabolic flux analysis was performed by assessing ^13^C-labeling of TCA intermediates in the Advanced Technology Core Facility at the Baylor College of Medicine using published protocols (Vantaku et al., 2017; Jin et al., 2017; Piyarathna et al., 2018; Kornberg et al., 2018), building on prior studies (Buescher et al., 2015; Butterfield and Halliwell, 2019; Jang et al., 2018; Lewis et al., 2014; Ruiz et al., 2015). In brief homogenized samples were extracted in chloroform/methanol/water, and analysis was performed on a 6490 QQQ triple quadrupole mass spectrometer (Agilent Technologies) coupled to a 1290 Series HPLC system via selected reaction monitoring (SRM). Metabolites were targeted in both positive and negative ion modes with an electrospray source ionization (ESI) voltage of +4,000 V in the positive ion mode and – 3,500 V in the negative ion mode. For each detected metabolite, ∼9-12 data points were collected. For the calculation of labeled metabolites, we used the equation (Mi*i)/n = Mi, the ratio of ^13^C-labeled metabolite to the total pool, where i = M+x (i.e., M+2, M+3, etc., values) and n = number of carbons in the respective metabolite (e.g., for citrate = 6). M+x values can be summed to yield the molar percent enrichment (MPE), with molar enrichment (ME) equal to MPE times the total pool mean.

#### Kinetic modeling of ^13^C incorporation into TCA cycle intermediates

As we have described (Doulias et al., 2023) using tenets developed by Gupta et al. (2009), we constructed a semi-quantitative kinetic model for analysis of changes in metabolic flux through consecutive enzymatic reactions of the TCA cycle in response to changes in the S-nitrosylation status of the corresponding enzymes. For multistep segments of the pathway containing unmeasured metabolites, the consecutive reactions were reduced to a single step, containing only measured substrate and product, i.e., the segment between citrate and α-ketoglutarate was analyzed as if it were a single-step reaction catalyzed by one-step enzymatic entity (that combines the activities of Aco and IDH since isocitrate was not measured in the mass spectrometric analysis). Similarly, a two-step conversion of α-ketoglutarate to succinate was considered as if a single step. The reaction rates (fluxes) were considered to follow mass-action law kinetics, consistent with the assumption that the substrate concentrations were much smaller in comparison to the corresponding Michaelis constant *K*_m_ for enzymatic reactions for the first turn through the TCA cycle. Accordingly, the flux through each step of the TCA cycle was assumed to be a product of substrate concentration and a corresponding kinetic constant and represent a pseudo first-order reaction process. These assumptions closely follow those utilized in the study by Gupta et al. (2009).

Citrate synthase reaction represents an entry point into the TCA cycle for two-carbon moieties in the form of acetyl-CoA (Ac-CoA), and is, as such, a two-substrate reaction. We presumed a constant level of Ac-CoA, as determined by a steady supply of carbons from a virtually unlimited pool of exogenously added lactate (10 mM in the bulk medium). Thus, the flux of the two-substrate reaction can be reduced to pseudo-first order form with observable kinetic constant k_1_ equal to a product of two constants, k_1_ = k_1_’ × [Ac-CoA], where k_1_’ is the actual kinetic constant for the two-substrate reaction.

Therefore, our model can be described by six elementary fluxes J_i_= k_i_ × [A_i_], corresponding to the six measured metabolites (citrate, α-ketoglutarate, succinate, fumarate, malate, and oxaloacetate). Here, [A_i_] is the concentration of analyte number “i”, and k_i_ is the kinetic constant for reaction of A_i_, with i = 1, 2, …6. For each A_i_, the rate of change is d[A_i_]/dt= J_i-1_- J_i_. For example, malate is the fifth metabolite in the sequence, so the change in malate level d[malate]/dt= J_4_ - J_5_= k_4_ × [fumarate] – k_5_ × [malate]; in other words, change in malate level equals the difference between malate production from fumarate and malate consumption in malate dehydrogenase reaction.

The initial pools of metabolites were set by a prolonged 6-hour starvation period when no source of two-carbon moieties (i.e., sucrose, pyruvate, nor lactate) were provided to the cells. The assumption was made under these conditions that all TCA cycle metabolites were converted to oxaloacetate and cannot be further converted due to lack of the second substrate of the citrate synthase reaction (Ac-CoA). Therefore, the initial condition for simulation was that the level of oxaloacetate equaled the total pool of TCA cycle metabolites, and all other metabolite levels were set to zero; all metabolites were unlabeled prior to the addition of ^13^C-labeled lactate. Throughout the period of addition, all the entering two-carbon moieties were considered fully labeled, a reasonable assumption given such a high concentration (10 mM) of labeled lactate was provided.

Simulation of non-steady state kinetics was performed computationally. The model was applied recursively, starting from the initial conditions, and changing the levels of each metabolite at each recursive step by an increment Δ[A_i_]_i_ ≈ d[A_i_]/dt to generate approximate theoretical kinetic curves for isotopologues of each metabolite. This approximation is valid as long as the time increment corresponding to a recursion step is sufficiently small compared to the characteristic time of the total processes. As we were interested only in relative changes upon the biological perturbation triggered by the PS1 mutation in the AD-hiN in the presence and absence of protein S-nitrosylation (the absence induced by pre-treatment with L-NAME), we were free to select an arbitrary time scale that satisfies this assumption. Moreover, the control simulation implied perfectly balanced fluxes throughout all steps. For evaluation of responses to perturbation, the parameters of the model were manually fitted to produce changes in the ratios of consecutive metabolites ([A_i-1_]/[A_i_]) within 3% error of experimentally observed values upon challenge related to nitrosative stress.

#### Bioenergetics assays on hiN with the Seahorse flux analyzer

Bioenergetic status of hiN was assessed using Seahorse XF^e^96 flux analyzer. Human neural progenitor cells were differentiated into hiN in the 60 inner wells of a 96-well Seahorse cell culture plate for 5-6 weeks at a density of 4×10^4^ cells per well (consisting of pure neurons without glial cells). At the start of the experiment, the differentiation medium was replaced with 190 µL assay medium supplemented in treatment well with dimethyl succinate (DMS, 5 mM, Acros). The basal assay medium was custom ordered Neurobasal medium (lacking sodium bicarbonate, HEPES, glucose, pyruvate and Phenol Red); it was supplemented with 10 mM glucose, 20 mM sodium pyruvate, 2 mM GlutaMAX, and 5 mM HEPES-sodium, pH 7.4. For the mitochondrial stress test assay, replicate readings of basal respiration OCR were followed by injections of ∼10 µL of 10x mitochondrial inhibitors to induce specific metabolic states of mitochondria as follows: The resting state (State 4) was induced by addition of 2 µg/ml Oligomycin; maximal respiration (State 3u) was induced by two sequential additions of 90-120 µM uncoupler DNP; and the experiment was terminated by addition of 2 µM of the respiratory inhibitor myxothiazol. Parallel measurements of ECAR were used to estimate the glycolytic contribution to total energy turnover of the cells. After the assay, cells were fixed with 4% paraformaldehyde for 15 minutes and stained with Hoechst 33342 (1:500) for subsequent cell counts. For normalization of bioenergetic parameters to cell number, hiN were counted using an ImageXpress Micro Confocal platform, quantifying the central area of each well (inside the ‘posts’) that contributes to Seahorse-measured fluxes, and then OCR readings were divided by cell numbers for each well.

#### Assessment of ATP production and turnover in AD-hiN and WT/Control hiN

Cellular ATP demand and rates of ATP production were estimated essentially as published previously (Andreyev et al., 2020) from the OCR and ECAR data using the known stoichiometry between ATP and oxygen and lactate, respectively. ECAR rates expressed in mpH/min reported by the Seahorse XF^e^96 flux analyzer were converted to proton production rate (PPR) expressed as [H^+^]/min. The conversion factor for this was determined by titrating the assay medium, the buffering capacity of which is provided predominantly by the 5 mM HEPES in the medium, with sequential pulses of sulfuric acid (data not shown). The pH response was virtually linear over the working range (6.8 – 7.4) and yielded a conversion factor of 6.07 pmol H^+^ per 1 mpH.

ECAR includes contributions from both lactic acid and CO_2_ in the form of carbonic acid. To determine the activity of glycolytic (lactate-producing) pathway, acidification due to CO_2_ produced by respiration has to be subtracted from the total ECAR value. The factor relating change in acidification to respiratory activity was determined empirically by dividing the decrease in ECAR by the drop in OCR upon inhibition of respiration with myxothiazol at the end of the Seahorse run. The product of this factor and OCR was subtracted from ECAR in the basal and/or maximally uncoupled states. The differences, corresponding to glycolytic activities, were converted to PPR using the factor mentioned above; PPR is equivalent to lactate production rate (1:1 lactate to H^+^ given almost complete dissociation at pH 7.4). Since one molecule of ATP is produced per each lactate molecule (1:1 ATP to lactate) in the glycolytic pathway, the rate of ATP production assumes the same value. For ATP production during oxidative phosphorylation, OCR was multiplied by 6, corresponding to the known stoichiometry of 36 ATP per glucose. Both rates were summed up to yield total ATP production rate.

According to the concept of respiratory control rate of ATP production, in the basal state this is equal to demand for ATP by the cells; under normal circumstances, this value would be less than the capacity to produce ATP. The capacity to produce ATP by oxidative phosphorylation or by glycolysis was calculated from the maximal OCR (State 3u) and from the ECAR value in the respiratory-inhibited cells (determined after addition of myxothiazol), respectively.

#### Analysis of the NAD^+^/NADH ratio in AD-hiN and WT/Control-hiN

Redox status of pyridine nucleotides (e.g., NADH/NAD^+^) in hiN cultures was assessed using autofluorescence (λ_ex_ = 340, λ_em_ = 460 nm) employing a Zeiss Axiovert100 microscope equipped with a Lambda DG4 (Sutter Instruments) light source (Kushnareva et al., 2013). The excitation and emission filters were 340/26 and 470/40 nm (maximal wavelength and bandwidth), respectively. Tetramethylrhodamine, ethyl ester (TMRE, 10 nM) was used as a mitochondrial counterstain (with 560/55 excitation and 655/40 emission). Images were acquired using MetaMorph software (Molecular Devices) and analyzed using Fiji (ImageJ) freeware. Punctate objects representing mitochondria were identified using Find Maxima function and the intensity of the brightest 200 of them (starting from the 6^th^ brightest to avoid bright artifacts) was measured. In addition to monitoring the NADH/NAD^+^ ratio at the basal state, reflecting steady-state conditions of active NADH utilization for ATP synthesis, the resting state induced by 2 µg/ml oligomycin was analyzed. In the resting state NADH utilization for ATP synthesis is blocked, and the reduced level of pyridine nucleotides are maximized.

#### Cell-based quantification of synapses with ImageXpress Micro Confocal high-content imaging system

For detection of synapses in AD-hiN and WT/Control hiN, their respective neural progenitor cells were differentiated into cerebrocortical neurons (with no glia) in the 60 inner wells of a 96-well plate (Corning, #3603) for 5-7 weeks at a density of ∼4×10^4^ cells per well (similar to the Seahorse experimental protocol). The cells were treated with 5 mM DMS or vehicle medium for 48 hours prior to assessment (10 wells per genotype per treatment were analyzed). hiN were then fixed with 4% paraformaldehyde for 15 min, permeabilized and blocked with blocking buffer (PBS supplemented with 3% BSA and 0.3% Triton X-100) for 30 min. The cells were decorated with primary antibodies for Synapsin 1 (rabbit monoclonal, Cell Signaling #5297S, 1:500), Homer 1 (mouse monoclonal, Synaptic Systems #160011, 1:250), and MAP2 (chicken polyclonal, Invitrogen #PA-10005, 1:500) in the blocking buffer overnight. Cells were washed with PBS and incubated with anti-rabbit AlexaFluor Plus 647 (Sigma A32795), anti-mouse AlexaFluor Plus 555 (Sigma A32773), anti-chicken AlexaFluor Plus 488 (Sigma A32931) secondary antibodies (1:500 in the blocking buffer) for 2 hours. The cells were also stained with Hoechst 33342 (1:500) to label nuclei for cell counts.

Images were acquired at 20x magnification in the ImageXpress Micro Confocal platform and analyzed using MetaXpress software (Molecular Dynamics, Inc.) equipped with a custom neurite and synapse detection and quantification module. For synapse detection, we identified pre- and postsynaptic terminals in close apposition (∼0.3 µm) (Trotter et al., 2019; Guo et al., 2019; Jiang et al., 2020; Zhou et al., 2022). Pre- and postsynaptic terminals were identified as Synapsin 1 and Homer 1-positive punctae, respectively. Neuronal bodies and neurites were identified by staining for MAP2. Coincidence with MAP2 stain was assessed to ensure the synaptic punctae were associated with neurons. The number of nuclei was counted separately with Hoechst staining and presumed to correspond to cell number. For each well, we calculated the number of synapses per cell. This number was averaged over each 10-well experimental group for each plate.

### QUANTIFICATION AND STATISTICAL ANALYSIS

Data were analyzed with the tests indicated in the appropriate paragraph for statistical analysis and bioinformatics (see above). For S-nitrosylation experiments and metabolic flux data, numbers (n) of biological replicates are included in the figure legends; comparisons were made by ANOVA followed by a PLSD post hoc test (Frane, 2019). For both the analysis of S-nitrosylated proteins and metabolic flux data, a p-value < 0.05 was considered significant.

#### Data availability

Source data are provided with this paper. The MS raw data have been deposited in the ProteomeXchange Consortium (http://proteomecentral.proteomexchange.org) via the PRIDE partner repository with the dataset identifier PXD042436.

## SUPPLEMENTAL INFORMATION

Supplemental information can be found online at https://doi.org/xxxxxxxx

## ACKNOWLEDGMENTS

We are indebted to Christian Metallo (Salk Institute), Itzhak Nissim (University of Pennsylvania), and John Essigmann (MIT) for helpful discussions and advice on the analysis of the metabolic flux results. We gratefully acknowledge access to the Baylor College of Medicine Advanced Technology Core directed by Nagireddy Putluri and thank Vasanta Putluri, who ran our metabolic flux experiments. We thank Eliezer Masliah (UC San Diego/NIA) and Robert Rissman (UC San Diego) for providing human brain tissues as part of the NIH supported National AD/ADRD Brain Bank. We thank Hossein Fazelinia, Ding Hua, and Lynn Spruce at the Proteomics Core Facility of the Children’s Hospital of Philadelphia Research Institute for their support and discussions regarding the MS-based proteomic analyses. This study was supported in part by the National Institutes of Health (NIH) grants R01 AG061845, R61 NS122098, and RF1 NS123298 (to T.N.), and R35 AG071734, RF1 AG057409, R56 AG065372, R01 DA048882 and DP1 DA041722 (to S.A.L.) and R01 AG056259 (to S.A.L. and S.R.T.). H.I. is the Gisela and Dennis Alter Research Professor at the Children’s Hospital of Philadelphia Research Institute.

## AUTHOR CONTRIBUTIONS

Conceptualization, S.A.L.; methodology, A.A., P.-T.D., H. Y., N.D., S.R.T., H.I., and S.A.L.; investigation, A.A., P.-T.D., H. Y., N.D., X.Z., M.L., M.B., C.B., and I.V.; visualization, P.- T.D., H. Y., N.D., A.A., X.Z., and T.N.; funding acquisition, T.N., S.R.T., H.I., and S.A.L.; writing – original draft, S.A.L.; writing – review & editing, P.-T.D., H.Y., N.D., A.A., X.Z., T.N., S.R.T., H.I., and S.A.L.

## REFERENCES

Akhtar, M.W., Sanz-Blasco, S., Dolatabadi, N., Parker, J., Chon, K., Lee, M.S., Soussou, W., McKercher, S.R., Ambasudhan, R., Nakamura, T., et al. (2016). Elevated glucose and oligomeric β-amyloid disrupt synapses via a common pathway of aberrant protein S-nitrosylation. Nat. Commun. 7, 10242.

Almeida, A., Almeida, J., Bolanos, J.P., Moncada, S. (2001). Different responses of astrocytes and neurons to nitric oxide: the role of glycolytically generated ATP in astrocyte protection. Proc. Natl. Acad. Sci. USA 98, 15294–15299.

Andreyev, A.Y., Tsui, H.S., Milne, G.L., Shmanai, V.V., Bekish, A.V., Fomich, M.A., Pham, M. N., Nong, Y., Murphy, A.N., Clarke, C. F., and Shchepinov, M. S. (2015). Isotope-reinforced polyunsaturated fatty acids protect mitochondria from oxidative stress. Free Rad. Biol. Med. 82, 63–72.

Andreyev, A. Y., Kushnareva, Y. E., Starkova, N. N. & Starkov, A. A. Metabolic ROS Signaling: To Immunity and Beyond. Biochemistry (Mosc) 85, 1650–1667 (2020).

Antunesm, F., Boveris, A., Cadenas, E (2004). On the mechanism and biology of cytochrome oxidase inhibition by nitric oxide. Proc. Natl. Acad. Sci. USA 101, 16774–16779.

Arnold, P.K., and Finley, L.W.S. (2022). Regulation and function of the mammalian tricarboxylic acid cycle. J. Biol. Chem. 299, 102838.

Barsoum, M.J., Yuan, H., Gerencser, A.A., Liot, G., Kushnareva, Y., Graber, S., Kovacs, I., Lee, W.D., Waggoner, J., Cui, J., et al. (2006). Nitric oxide-induced mitochondrial fission is regulated by dynamin-related GTPases in neurons. EMBO J. 25, 3900–3911.

Beltran, B., Orsi, A., Clementi, E., Moncada, S. (2000). Oxidative stress and S-nitrosylation of proteins in cells. Br. J. Pharmacol. 129, 953–960.

Bhaduri, A., Andrews, M.G., Kriegstein, A.R., and Nowakowski, T.J. (2020). Are organoids ready for prime time? Cell Stem Cell 27, 361–365.

Bolanos, J.P., Peuchen, S., Heales, S.J., Land, J.M., Clark, J.B. (1994). Nitric oxide-mediated inhibition of the mitochondrial respiratory chain in cultured astrocytes. J. Neurochem. 63, 910– 916.

Brown, G.C., and Cooper, C.E. (1994). Nanomolar concentrations of nitric oxide reversibly inhibit synaptosomal respiration by competing with oxygen at cytochrome oxidase. FEBS Lett. 356, 295–298.

Bruegger, J.J., Smith, B.C., Wynia-Smith, S.L., and Marletta, M.A. (2018). Comparative and integrative metabolomics reveal that S-nitrosation inhibits physiologically relevant metabolic enzymes. J. Biol. Chem. 293, 6282–6296.

Buescher, J.M., Antoniewicz, M.R., Boros, L.G., Burgess, S.C., Brunengraber, H., Clish, C.B., DeBerardinis, R.J., Feron, O., Frezza, C., Ghesquiere, B., et al. (2015). A roadmap for interpreting (13)C metabolite labeling patterns from cells. Curr. Opin. Biotechnol. 34, 189–201.

Butterfield, D.A., and Halliwell, B. (2019). Oxidative stress, dysfunctional glucose metabolism and Alzheimer disease. Nat. Rev. Neurosci. 20, 148–160.

Caminiti, S.P., Sala, A., Iaccarino, L., Beretta, L., Pilotto, A., Iannaccone, S., Magnani, G. Padovani, A. Refini-Strambi, L., and Perani, D. (2019) Brain glucose metabolism in Lewy body dementia: implications for diagnostic criteria. Alz. Res. Therapy 11, 20.

Cheng, X.T., Huang, N., and Sheng, Z.H. (2022) Programming axonal mitochondrial maintenance and bioenergetics in neurodegeneration and regeneration. Neuron 110, 1899–1923.

Cho, D.H., Nakamura, T., Fang, J., Cieplak, P., Godzik, A., Gu, Z., and Lipton, S.A. (2009). S-Nitrosylation of Drp1 mediates β-amyloid-related mitochondrial fission and neuronal injury. Science 324, 102–105.

Chung, C.Y., Khurana, V., Auluck, P.K., Tardiff, D.F., Mazzulli, J.R., Soldner, F., Baru, V., Lou, Y., Freyzon, Y., Cho, S., et al. (2013). Identification and rescue of α-synuclein toxicity in Parkinson patient-derived neurons. Science 342, 983–987.

Clementi, E., Brown, G.C., Feelisch, M., Moncada, S. (1998). Persistent inhibition of cell respiration by nitric oxide: crucial role of S-nitrosylation of mitochondrial complex I and protective action of glutathione. Proc. Natl. Acad. Sci. USA 95, 7631–7636.

Cooper, C.E., and Giulivi, C. (2007). Nitric oxide regulation of mitochondrial oxygen consumption II: Molecular mechanism and tissue physiology. Am. J. Physiol. Cell Physiol. 292, C1993–2003

Cooper, O., Seo, H., Andrabi, S., Guardia-Laguarta, C., Graziotto, J., Sundberg, M., McLean, J.R., Carrillo-Reid, L., Xie, Z., Osborn, T., et al. (2012). Pharmacological rescue of mitochondrial deficits in iPSC-derived neural cells from patients with familial Parkinson’s disease. Sci. Transl. Med. 4, 141ra190.

Cordes, T., and Metallo, C.M. (2019). Quantifying intermediary metabolism and lipogenesis in cultured mammalian cells using stable isotope tracing and mass spectrometry. Methods Mol. Biol. 1978, 219–241.

Chouchani, E.T. et al. (2014). Ischaemic accumulation of succinate controls reperfusion injury through mitochondrial ROS. Nature 515, 431–435.

Czaniecki, C., Ryan, T., Stykel, M.G., Drolet, J., Heide, J., Hallam, R., Wood, S., Coackley, C., Sherriff, K., Bailey, C.D.C., et al. (2019). Axonal pathology in hPSC-based models of Parkinson’s disease results from loss of Nrf2 transcriptional activity at the Map1b gene locus. Proc. Natl. Acad. Sci. U. S. A. 116, 14280–14289.

DeKosky, S.T., and Scheff, S.W. (1990). Synapse loss in frontal cortex biopsies in Alzheimer’s disease: correlation with cognitive severity. Ann Neurol. 27, 457–64.

Dembitskaya, Y., Piette, C., Perez, S., Berry, H., Magistretti, P. J., and Venance, L. (2022). Lactate supply overtakes glucose when neural computational and cognitive loads scale up. Proc. Natl. Acad. Sci. U. S. A. 119, e2212004119.

Diaz-Garcia, C.M., Mongeon, R., Lahmann, C., Koveal, D., Zucker, H., and Yellen, G. (2017). Neuronal stimulation triggers neuronal glycolysis and not lactate uptake. Cell Metab. 26, 361–374 e364.

Doulias, P.T., Greene, J.L., Greco, T.M., Tenopoulou, M., Seeholzer, S.H., Dunbrack, R.L., and Ischiropoulos, H. (2010). Structural profiling of endogenous S-nitrosocysteine residues reveals unique features that accommodate diverse mechanisms for protein S-nitrosylation. Proc. Natl. Acad. Sci. U. S. A. 107, 16958–16963.

Doulias, P.T., Nakamura, T., Scott, H., McKercher, S.R., Sultan, A., Deal, A., Albertolle, M., Ischiropoulos, H., and Lipton, S.A. (2021). TCA cycle metabolic compromise due to an aberrant S-nitrosoproteome in HIV-associated neurocognitive disorder with methamphetamine use. J. Neurovirol. 27, 367–378.

Doulias, P.T., Tenopoulou, M., Greene, J.L., Raju, K., and Ischiropoulos, H. (2013a). Nitric oxide regulates mitochondrial fatty acid metabolism through reversible protein S-nitrosylation. Sci. Signal. 6, rs1.

Doulias, P.T., Tenopoulou, M., Raju, K., Spruce, L.A., Seeholzer, S.H., and Ischiropoulos, H. (2013b). Site specific identification of endogenous S-nitrosocysteine proteomes. J. Proteomics 92, 195–203.

Doulias, P.T., Yang, H. Andreyev, A., Dolatabadi, N., Scott, H., Raspur, C.K., Patel, P.R., Nakamura, T., Tannenbaum, S.R., Ischiropoulos, H., and Lipton, S.A. S-Nitrosylation-mediated dysfunction of TCA cycle enzymes in synucleinopathy studied in postmortem human brains and hiPSC-derived neurons. Cell Chem. Biol. 2023, in press.

Eckert, A. Schmitt, K., and Götz (2011). Mitochondrial dysfunction - the beginning of the end in Alzheimer’s disease? Separate and synergistic modes of tau and amyloid-β toxicity. J. Alzheimers Res Ther. 3, 15.

Exner, N., Lutz, A.K., Haass, C., and Winklhofer, K.F. (2012). Mitochondrial dysfunction in Parkinson’s disease: molecular mechanisms and pathophysiological consequences. EMBO J. 31, 3038–3062.

Fernandez-Garcia, J., Altea-Manzano, P., Pranzini, E., and Fendt, S.M. (2020). Stable isotopes for tracing mammalian-cell metabolism *in vivo*. Trends Biochem. Sci. 45, 185–201.

Frane, A. Type I Error Control in Psychology Research: Improving Understanding in General and Addressing Multiplicity in Some Specific Contexts. Thesis dissertation, UCLA, 2019. https://escholarship.org/content/qt8sn0z5g0/qt8sn0z5g0.pdf?t=pw82wy.

Gibson, G.E., Starkov, A., Blass, J.P., Ratan, R.R., and Beal, M.F. (2010). Cause and consequence: mitochondrial dysfunction initiates and propagates neuronal dysfunction, neuronal death and behavioral abnormalities in age-associated neurodegenerative diseases. Biochim. Biophys. Acta 1802, 122–134.

Ghatak, S., Dolatabadi, N., Trudler, D., Zhang, X.T., Wu, Y., Mohata, M., Ambasudhan, R., Talantova, M., and Lipton, S.A. (2019). Mechanisms of hyperexcitability in Alzheimer’s disease hiPSC-derived neurons and cerebral organoids vs isogenic controls. eLife 9, 8.

Ghatak, S., Dolatabadi, N., Gao, R., Wu, Y., Scott, H., Trudler, D., Sultan, A., Ambasudhan, R., Nakamura, T., Masliah, E., Talantova, M., Voytek, B., and Lipton, S.A. (2021). NitroSynapsin ameliorates hypersynchronous neural network activity in Alzheimer hiPSC models. Mol. Psychiatry 26, 5751–5765.

González-Rodriguez, P., Zampese, E., Stout, K.A., Guzman, J.N., Ilijic, E., Yang, B., Tkatch, T., Stavarache, M.A., Wokosin, D.L., Gao, L., et al. (2021). Disruption of mitochondrial complex I induces progressive parkinsonism. Nature 599, 650–656.

Guo, S. M. et al. (2019). Multiplexed and high-throughput neuronal fluorescence imaging with diffusible probes. Nat Commun 10, 4377.

Gupta, S., Maurya, M.R., Stephens, D.L., Dennis, E.A., and Subramaniam, S. (2009). An integrated model of eicosanoid metabolism and signaling based on lipidomics flux analysis. Biophys. J. 96, 4542–4551.

Jang, C., Chen, L., and Rabinowitz, J.D. (2018). Metabolomics and isotope tracing. Cell 173, 822–837.

Jiang, H. et al. (2020). Live Neuron High-Content Screening Reveals Synaptotoxic Activity in Alzheimer Mouse Model Homogenates. Sci Rep 10, 3412.

Jin, F., Thaiparambil, J., Donepudi, S. R., Vantaku, V., Piyarathna, D., Maity, S., Krishnapuram, R., Putluri, V., Gu, F., Purwaha, P., Bhowmik, S. K., Ambati, C. R., von Rundstedt, F. C., Roghmann, F., Berg, S., Noldus, J., Rajapakshe, K., Gödde, D., Roth, S., Störkel, S., Degener, S., Michailidis, G., Kaipparettu, B.A., Karanam, B., Terris, M.K., Kavuri, S.M., Lerner, S.P., Kheradmand, F., Coarfa, C., Sreekumar, A., Lotan, Y., El-Zein, R., Putluri, N. (2017). Tobacco-specific carcinogens induce hypermethylation, DNA adducts, and DNA damage in bladder cancer. Cancer Prev. Res. 10, 588–597.

Kam, T.I., Mao, X., Park, H., Chou, S.C., Karuppagounder, S.S., Umanah, G.E., Yun, S.P., Brahmachari, S., Panicker, N., Chen, R., et al. (2018). Poly(ADP-ribose) drives pathologic α-synuclein neurodegeneration in Parkinson’s disease. Science 362, eaat8407.

Kornberg, M.D., Bhargava, P., Kim, P.M., Putluri, V., Snowman, A.M., Putluri, N., Calabresi, P.A., Snyder, S. H. (2018). Dimethyl fumarate targets GAPDH and aerobic glycolysis to modulate immunity. Science 360, 449–453.

Kriks, S., Shim, J.W., Piao, J., Ganat, Y.M., Wakeman, D.R., Xie, Z., Carrillo-Reid, L., Auyeung, G., Antonacci, C., Buch, A., et al. (2011). Dopamine neurons derived from human ES cells efficiently engraft in animal models of Parkinson’s disease. Nature 480, 547–551.

Kushnareva, Y.E., Gerencser, A.A., Bossy, B., Ju, W.K., White, A.D., Waggoner, J., Ellisman, M.H., Perkins, G., Bossy-Wetzel, E. (2013). Loss of OPA1 disturbs cellular calcium homeostasis and sensitizes for excitotoxicity. Cell Death Differ. 20, 353–565.

Laranjinha, J., Nunes, C., Ledo, A., Lourenço, C., Rocha, B., Barbosa, R.M. (2021). The peculiar facets of nitric oxide as a cellular messenger: from disease-associated signaling to the regulation of brain bioenergetics and neurovascular coupling. Neurochem. Res. 46, 64–76.

Ledo, A., Barbosa, R., Cadenas, E., Laranjinha, J. (2010). Dynamic and interacting profiles of NO and O_2_ in rat hippocampal slices. Free Radic. Biol. Med. 48, 1044–1050.

Lewis, C.A., Parker, S.J., Fiske, B.P., McCloskey, D., Gui, D.Y., Green, C.R., Vokes, N.I., Feist, A.M., Vander Heiden, M.G., and Metallo, C.M. (2014). Tracing compartmentalized NADPH metabolism in the cytosol and mitochondria of mammalian cells. Mol. Cell 55, 253–263.

Li, S., and Sheng, Z.H. (2022). Energy matters: presynaptic metabolism and the maintenance of synaptic transmission. Nat. Rev. Neurosci. 23, 4–22.

Lin, T., Ambasudhan, R., Yuan, X., Li, W., Hilcove, S., Abujarour, R., Lin, X., Hahm, H.S., Hao, E., Hayek, A., et al. (2009). A chemical platform for improved induction of human iPSCs. Nat. Methods 6, 805–808.

Lipton, S.A., Rezaie, T., Nutter, A., Lopez, K.M., Parker, J., Kosaka, K., Satoh, T., McKercher, S. R., Masliah, E., and Nakanishi, N. (2016). Therapeutic advantage of pro-electrophilic drugs to activate the Nrf2/ARE pathway in Alzheimer’s disease models. Cell Death Dis. 7, e2499.

Liu, T., Zhang, M., Mukosera, G.T., Borchardt, D., Li, Q., Tipple, T.E., Ishtiaq Ahmed, A.S., Power, G.G., and Blood, A.B. (2019). L-NAME releases nitric oxide and potentiates subsequent nitroglycerin-mediated vasodilation. Redox Biol. 26, 101238.

Magistretti, P.J., and Allaman, I. (2015). A cellular perspective on brain energy metabolism and functional imaging. Neuron 86, 883–901.

Magistretti, P.J., and Allaman, I. (2018). Lactate in the brain: from metabolic end-product to signalling molecule. Nat. Rev. Neurosci. 19, 235–249.

Martinez-Ruiz, A., Cadenas, S., Lamas, S. (2011). Nitric oxide signaling: classical, less classical, and nonclassical mechanisms. Free Radic. Biol. Med. 51, 17–29.

Mi, H., Muruganujan, A., Casagrande, J.T., and Thomas, P.D. (2013). Large-scale gene function analysis with the PANTHER classification system. Nat. Protoc. 8, 1551–1566.

Nakamura, T., and Lipton, S.A. (2017). ’SNO’-storms compromise protein activity and mitochondrial metabolism in neurodegenerative disorders. Trends Endocrinol. Metab. 28, 879–892.

Nakamura, T., and Lipton, S.A. (2020). Nitric oxide-dependent protein post-translational modifications impair mitochondrial function and metabolism to contribute to neurodegenerative diseases. Antioxid. Redox Signal. 32, 817–833.

Nakamura, T., Oh, C.K., Liao, L., Zhang, X., Lopez, K.M., Gibbs, D., Deal, A.K., Scott, H.R., Spencer, B., Masliah, E., et al. (2021). Noncanonical transnitrosylation network contributes to synapse loss in Alzheimer’s disease. Science 371, eaaw0843.

Paquet, D., Kwart, D., Chen, A., Sproul, A., Jacob, S., Teo, S., Olsen, K.M., Gregg, A., Noggle, S., and Tessier-Lavigne, M. (2016). Efficient introduction of specific homozygous and heterozygous mutations using CRISPR/Cas9. Nature 533, 125–129,

Pellerin, L., and Magistretti, P.J. (1994). Glutamate uptake into astrocytes stimulates aerobic glycolysis: a mechanism coupling neuronal activity to glucose utilization. Proc. Natl. Acad. Sci. U. S. A. 91, 10625–10629.

Piyarathna, D., Rajendiran, T. M., Putluri, V., Vantaku, V., Soni, T., von Rundstedt, F. C., Donepudi, S. R., Jin, F., Maity, S., Ambati, C. R., Dong, J., Gödde, D., Roth, S., Störkel, S., Degener, S., Michailidis, G., Lerner, S. P., Pennathur, S., Lotan, Y., Coarfa, C., Sreekumar, A., Putluri, N. (2018). Distinct lipidomic landscapes associated with clinical stages of urothelial cancer of the bladder. Eur. Urol. Focus 17, 30107–30114.

Raju, K., Doulias, P.T., Evans, P., Krizman, E. N., Jackson, J. G., Horyn, O., Daikhin, Y., Nissim, I., Yudkoff, M., Nissim, I., Sharp, K. A., Robinson, M. B., and Ischiropoulos, H. (2015). Regulation of brain glutamate metabolism by nitric oxide and S-nitrosylation. Sci. Signal. 8, ra68.

Ruiz, M., Gelinas, R., Vaillant, F., Lauzier, B., and Des Rosiers, C. (2015). Metabolic tracing using stable isotope-labeled substrates and mass spectrometry in the perfused mouse heart. Methods Enzymol. 561, 107–147.

Ryan, S.D., Dolatabadi, N., Chan, S.F., Zhang, X., Akhtar, M.W., Parker, J., Soldner, F., Sunico, C.R., Nagar, S., Talantova, M., et al. (2013). Isogenic human iPSC Parkinson’s model shows nitrosative stress-induced dysfunction in MEF2-PGC1α transcription. Cell 155, 1351–1364.

Ryan, T., Bamm, V.V., Stykel, M.G., Coackley, C.L., Humphries, K.M., Jamieson-Williams, R., Ambasudhan, R., Mosser, D.D., Lipton, S.A., Harauz, G., et al. (2018). Cardiolipin exposure on the outer mitochondrial membrane modulates α-synuclein. Nat. Commun. 9, 817.

San Martin, A., Arce-Molina, R., Galaz, A., Perez-Guerra, G., Barros, L.F. (2017). Nanomolar nitric oxide concentrations quickly and reversibly modulate astrocytic energy metabolism. J. Biol. Chem. 292, 9432–9438.

Schweizer, M., and Richter, C. (1994). Nitric oxide potently and reversibly deenergizes mitochondria at low oxygen tension. Biochem. Biophys. Res. Commun. 204, 169–175.

Selak, M.A. et al. (2005). Succinate links TCA cycle dysfunction to oncogenesis by inhibiting HIF-alpha prolyl hydroxylase. Cancer Cell 7, 77–85.

Seneviratne, U., Godoy, L.C., Wishnok, J.S., Wogan, G.N., and Tannenbaum, S.R. (2013). Mechanism-based triarylphosphine-ester probes for capture of endogenous RSNOs. J. Am. Chem. Soc. 135, 7693–7704.

Seneviratne, U., Nott, A., Bhat, V.B., Ravindra, K.C., Wishnok, J.S., Tsai, L.H., and Tannenbaum, S.R. (2016). S-nitrosation of proteins relevant to Alzheimer’s disease during early stages of neurodegeneration. Proc. Natl. Acad. Sci. U. S. A. 113, 4152–4157.

Soldner, F., Laganiere, J., Cheng, A.W., Hockemeyer, D., Gao, Q., Alagappan, R., Khurana, V., Golbe, L.I., Myers, R.H., Lindquist, S., et al. (2011). Generation of isogenic pluripotent stem cells differing exclusively at two early onset Parkinson point mutations. Cell 146, 318–331.

Stewart, V.C., Heales, S.J. (2003). Nitric oxide-induced mitochondrial dysfunction: implications for neurodegeneration. Free Radic. Biol. Med. 34, 287–303.

Talantova, M., Sanz-Blasco, S., Zhang, X., Xia, P., Akhtar, M.W., Okamoto, S.-i., Dziewczapolski, G., Nakamura, T., Cao, G., Pratt, A.E., Kang, Y.-J., Tu, S., Molokanova, E., McKercher, S.R., Hires, A., Sason, H., Stouffer, D.G., Buczynski, M.W., Solomon, J., Michael, S., Powers, E.T., Kelly, J.W., Roberts, A.J., Tong, G., Fang-Newmeyer, T., Parker, J., Holland, E.A., Zhang, D., Nakanishi, N., Chen, H.-S.V., Wolosker, H., Parsons, L.H., Ambasudhan, R., Masliah, E., Heinemann, S.F., Piña-Crespo, J.C., Lipton, S.A. (2013). Aβ induces astrocytic glutamate release, extrasynaptic NMDA receptor activation, and synaptic loss. Proc. Natl. Acad. Sci. USA 110, E2518–2527.

Terry, R.D., Masliah, E., Salmon, D.P., Butters, N., DeTeresa, R., Hill, R., Hansen, L.A., Katzman, R. (1991). Physical basis of cognitive alterations in Alzheimer’s disease: synapse loss is the major correlate of cognitive impairment. Ann Neurol. 30, 572–580.

Torrini, C., Nguyen, T.T.T., Shu, C., Mela, A., Humala, N., Mahajan, A., Seeley, E.H., Zhang, G., Westhoff, M.A., Karpel-Massler, G., Bruce, J.N., Canoll, P., Siegelin, M.D. (2022). Lactate is an epigenetic metabolite that drives survival in model systems of glioblastoma. Mol. Cell S1097-2765(22)00647-5.

Traxler, L., Herdy, J. R., Stefanoni, D., Eichhorner, S., Pelucchi, S., Szücs, A., Santagostino, A., Kim, Y., Agarwal, R.K., Schlachetzki, J.C.M., Glass, C.K., Lagerwall, J., Galasko, D., Gage, F.H., D’Alessandro, A., and Mertens, J. (2022). Warburg-like metabolic transformation underlies neuronal degeneration in sporadic Alzheimer’s disease. Cell Metab. 34, 1248–1263.e6.

Trotter, J. H. et al. (2019). Synaptic neurexin-1 assembles into dynamically regulated active zone nanoclusters. J Cell Biol 218, 2677–2698.

Trudler, D., Sanz-Blasco, S., Eisele, Y.S., Ghatak, S., Bodhinathan, K., Akhtar, M.W., Lynch, W.P., Pina-Crespo, J.C., Talantova, M., Kelly, J.W., et al. (2021). α-Synuclein oligomers induce glutamate release from astrocytes and excessive extrasynaptic NMDAR activity in neurons, thus contributing to synapse loss. J. Neurosci. 41, 2264–2273.

Tzioras, M., McGeachan, R. I., Durrant, C. S., & Spires-Jones, T. L. (2023). Synaptic degeneration in Alzheimer disease. Nat. Rev. Neurol. 19, 19–38.

van Ooijen, M.P., Jong, V.L., Eijkemans, M.J.C., Heck, A.J.R., Andeweg, A.C., Binai, N.A., and van den Ham, H.J. (2018). Identification of differentially expressed peptides in high-throughput proteomics data. Brief Bioinform. 19, 971–981.

Vantaku, V., Donepudi, S.R., Ambati, C.R., Jin, F., Putluri, V., Nguyen, K., Rajapakshe, K., Coarfa, C., Battula, V.L., Lotan, Y., Putluri, N. (2017). Expression of ganglioside GD2, reprogram the lipid metabolism and EMT phenotype in bladder cancer. Oncotarget. 8, 95620– 95631.

Vazquez-Velez, G.E., and Zoghbi, H.Y. (2021). Parkinson’s disease genetics and pathophysiology. Annu. Rev. Neurosci. 44, 87–108.

Yang, H., Oh, C.-k., Amal, H., Wishnok, J.S., Lewis, S., Schahrer, E., Trudler, D., Nakamura, T., Tannenbaum, S.R., Lipton, S.A. (2022). Mechanistic insight into female predominance in Alzheimer’s disease based on aberrant protein S-nitrosylation of C3. Sci. Adv. 14;8:eade0764.

Zheng, X., Boyer, L., Jin, M., Mertens, J., Kim, Y., Ma, L., Ma, L., Hamm, M., Gage, F.H., and Hunter, T. (2016). Metabolic reprogramming during neuronal differentiation from aerobic glycolysis to neuronal oxidative phosphorylation. eLife 5, e13374.

Zhou, B., Lu, J. G., Siddu, A., Wernig, M. & Sudhof, T.C. (2022). Synaptogenic effect of APP-Swedish mutation in familial Alzheimer’s disease. Sci Transl Med 14, eabn9380.

