## Supplementary Information for "Metabolic bypass rescues aberrant S-nitrosylation-induced TCA cycle inhibition and synapse loss in Alzheimer’s disease human neurons"

**Supplemental Figures, Supplemental Figure Legends, and Supplemental Table Legends**

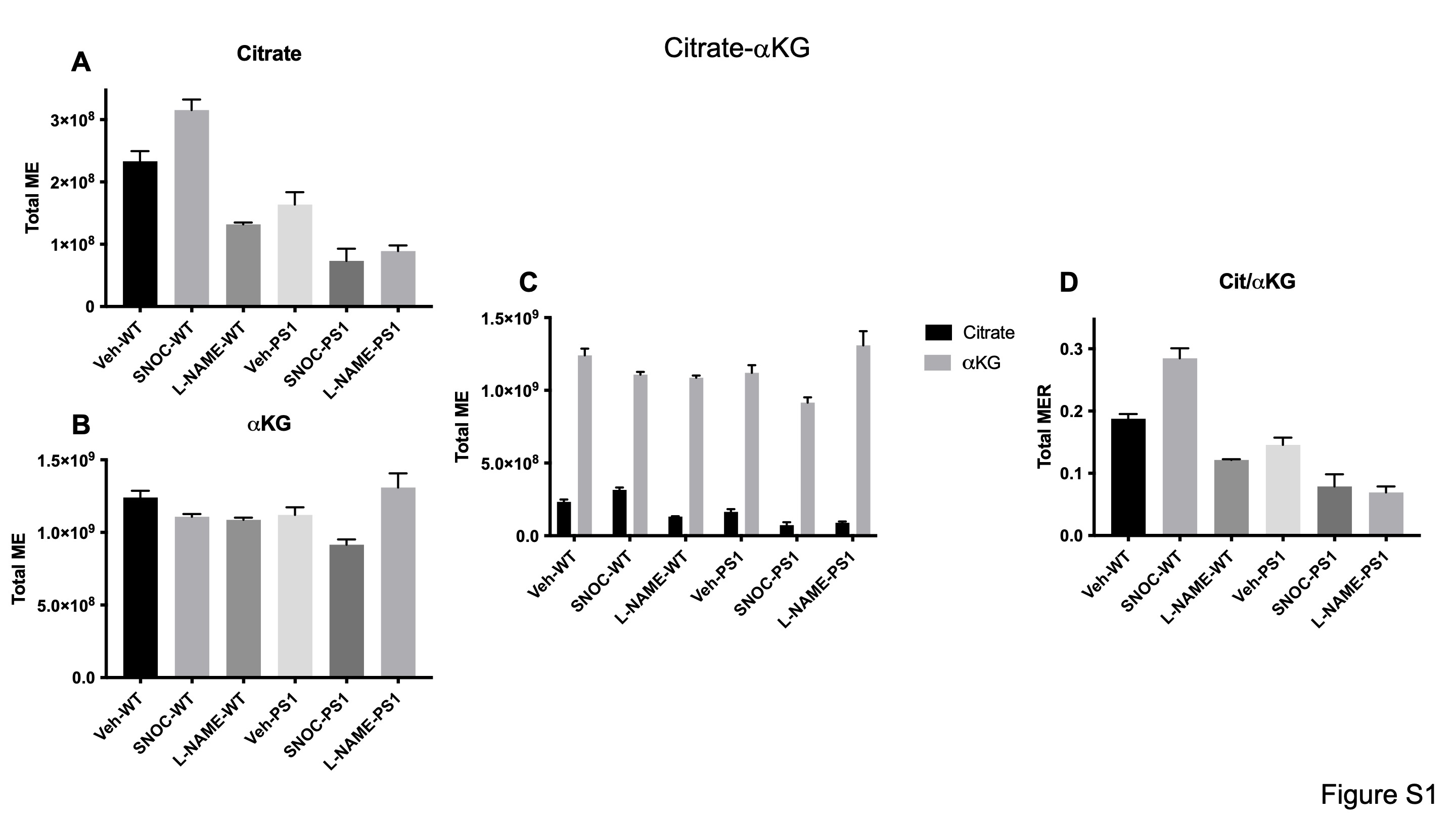

**Figure S1. Total ME and MER data for the pair of metabolites citrate and α-ketoglutarate (αKG), representing flux through aconitate and isocitrate dehydrogenase**

Values are mean + SEM (n = 3).

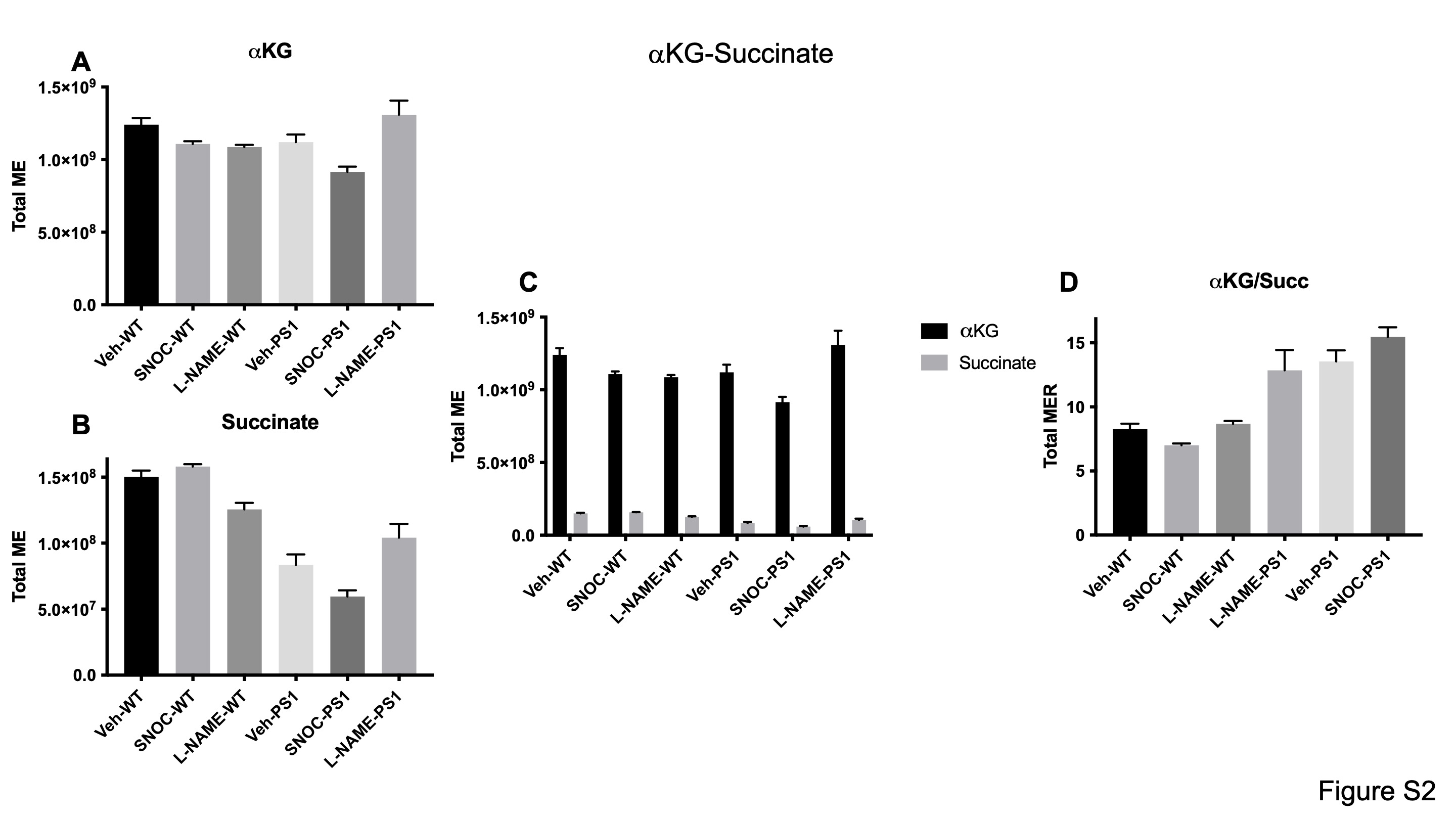

**Figure S2. Total ME and MER data for the pair of metabolites α-ketoglutarate (αKG) and succinate (Succ), representing flux through α-ketoglutarate dehydrogenase and succinyl coenzyme-A synthetase**

Values are mean + SEM (n = 3).

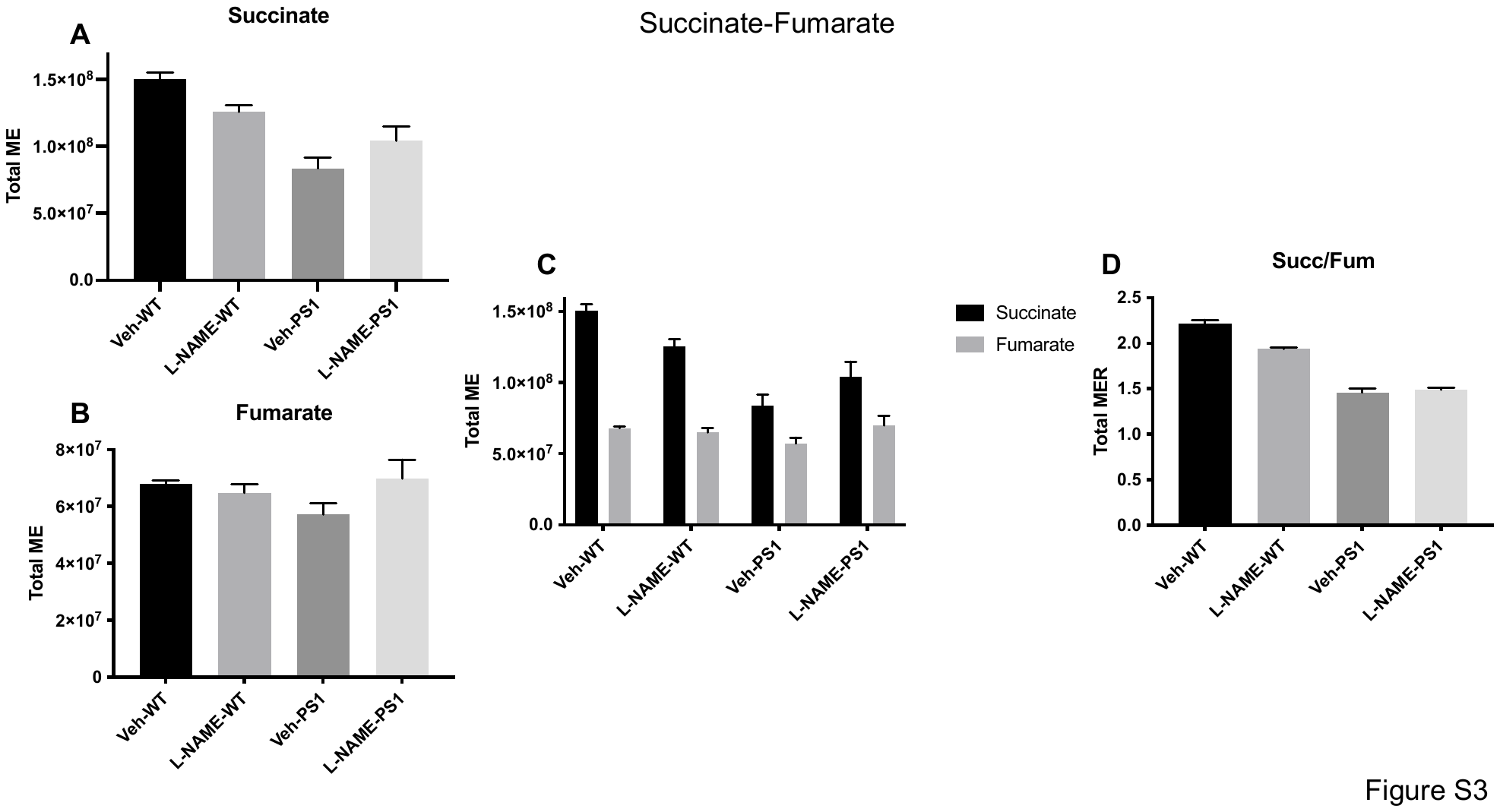

**Figure S3. Total ME and MER data for the pair of metabolites succinate (Succ) and fumarate (Fum), representing flux through succinate dehydrogenase**

Values are mean + SEM (n = 3).

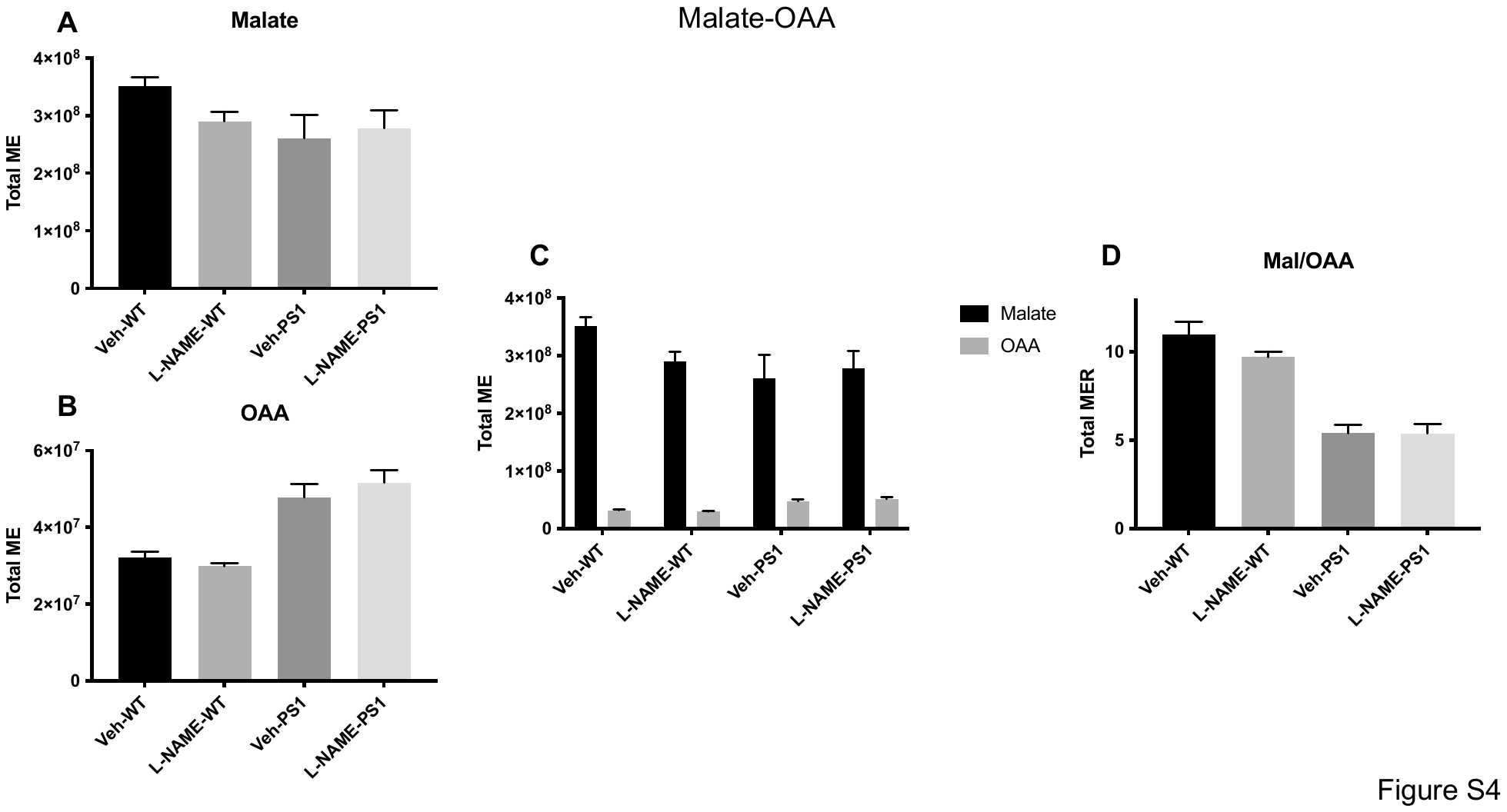

**Figure S4. Total ME and MER data for the pair of metabolites malate and oxaloacetate (OAA), representing flux through malate dehydrogenase**

Values are mean + SEM (n = 3).

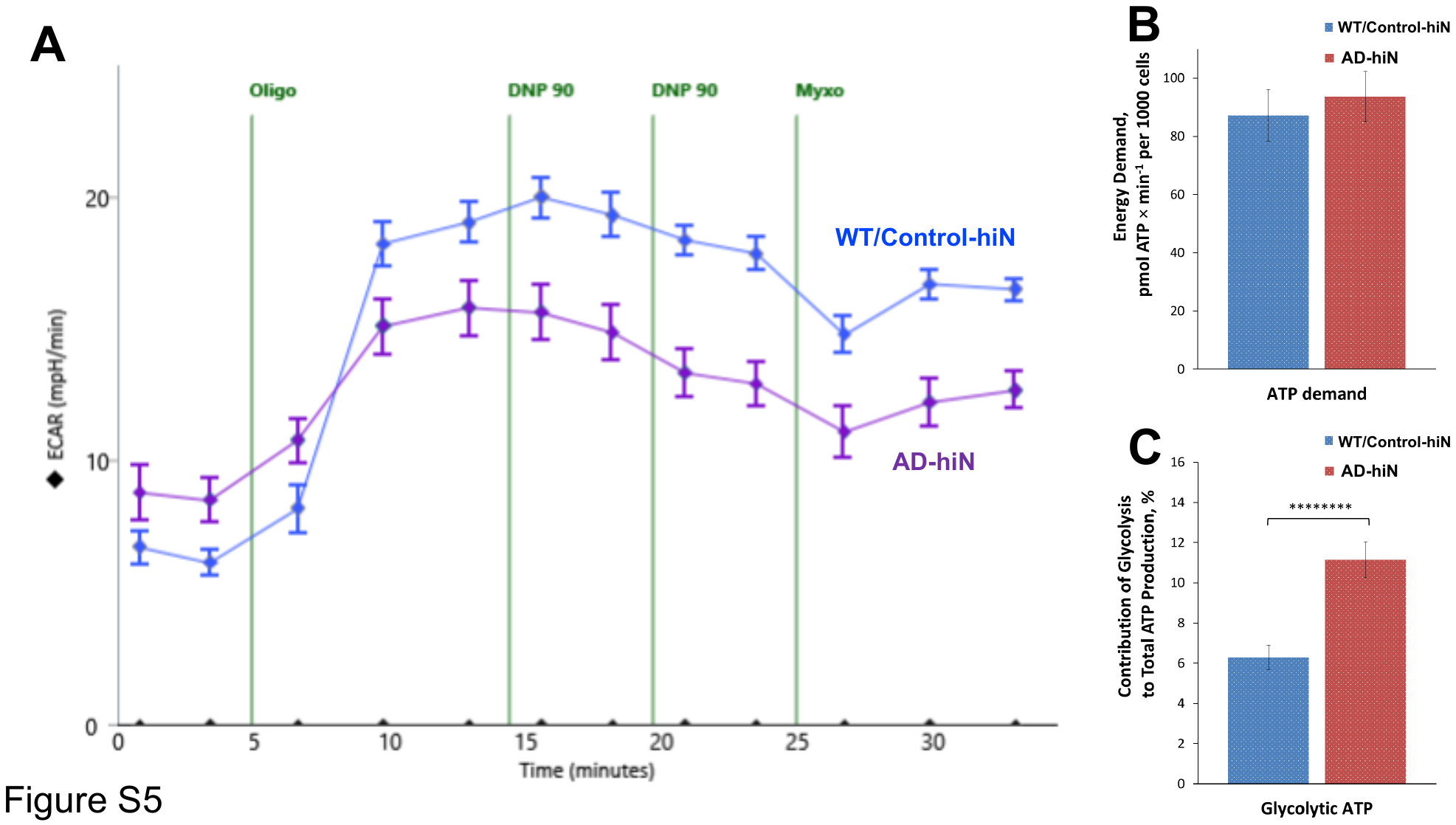

**Figure S5*.* Compensatory activation of aerobic glycolysis in respiratory-deficient AD-hiN (Warburg-like effect)**

(**A**) Representative experiment in Seahorse Flux Analyzer showing extracellular acidification rate (ECAR) for AD-hiN vs. WT/Control hiN. Injections are shown by vertical lines: Oligo, 2 μg/ml oligomycin; two injections of DNP, 90 μM; Myxo, 2 μM myxothiazol.

(**B, C**) Increased contribution of glycolysis to total ATP production in AD-hiN vs. WT/Control-hiN Total ATP production in the basal state of hiN that reflects cellular energy demand is unaffected by genotype (**B**). However, contribution of glycolysis to total ATP production increases about 2-fold in AD-hiN compared to WT/Control hiN (**C**), thus compensating in part for ATP observed during glycolysis in AD-hiN (see **Figure S6**). Data are mean± SEM; ***p < 0.001, by Student’s t-test (n = 22 plates analyzed, each in separate experiments for panel B; n = 43 plates for panel C).

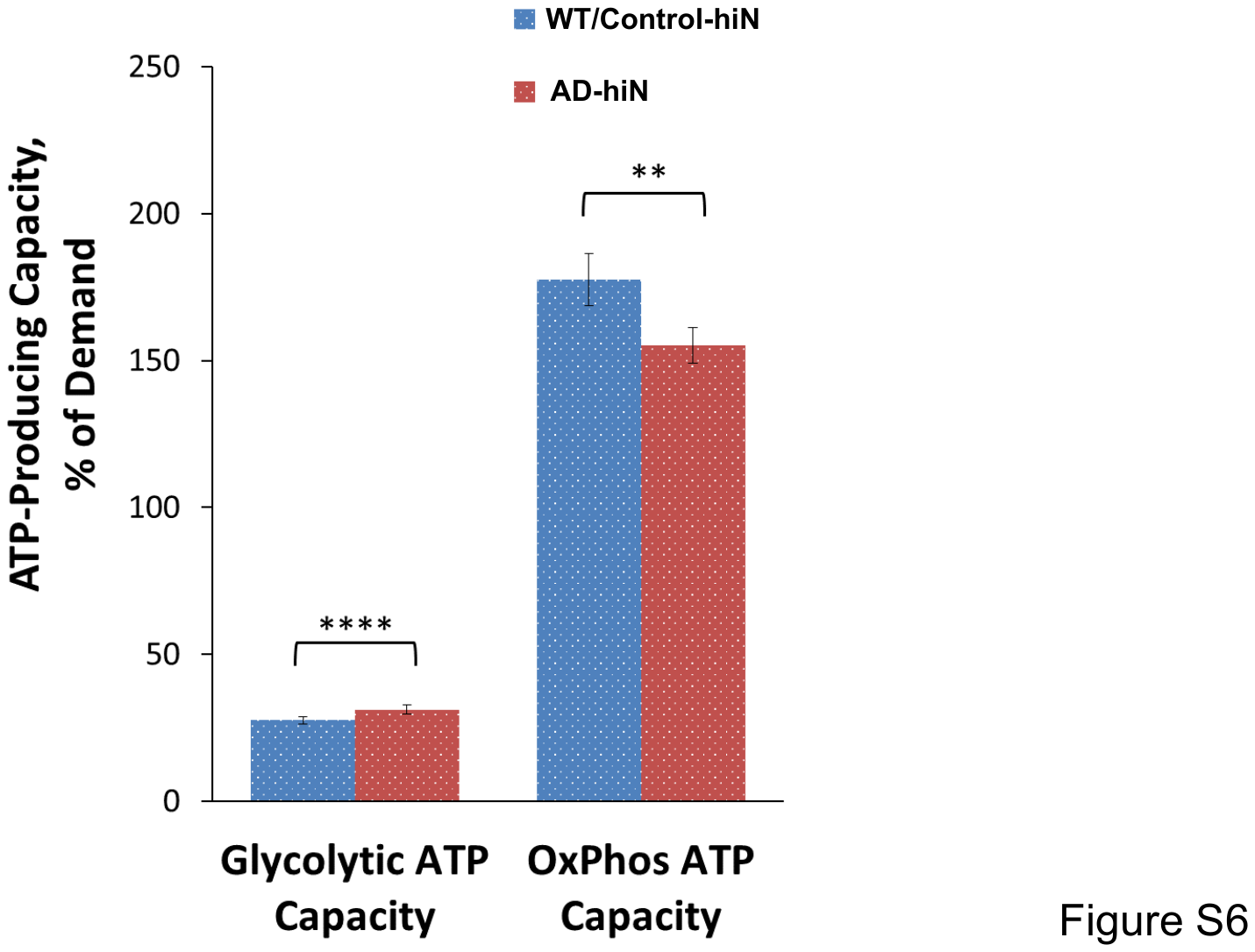

**Figure S6*.* Deficiency in spare energy-transducing capacity in AD-hiN neurons compared to WT/Control-hiN**

Calculated ATP-producing capacity (see METHOD DETAILS) for glycolysis (bars on *left*) and oxidative phosphorylation (OXPHOS, TCA cycle plus ETC) (bars on *right*) relative to total ATP demand (as shown in **Figure S5B**). Maximally-stimulated glycolysis by itself is incapable of meeting energy demands of hiN reaching only about 30-40% of required ATP production. OXPHOS possesses spare respiratory capacity in WT/Control-hiN which, however, is suppressed in AD-hiN (reaching ~170% in WT/Control-hiN but only ~150% in AD-hiN). Data are mean ± SEM; **p < 0.01, ****p < 0.0001 by Student’s t-test (n = 43 plates tested, each in a separate experiment).

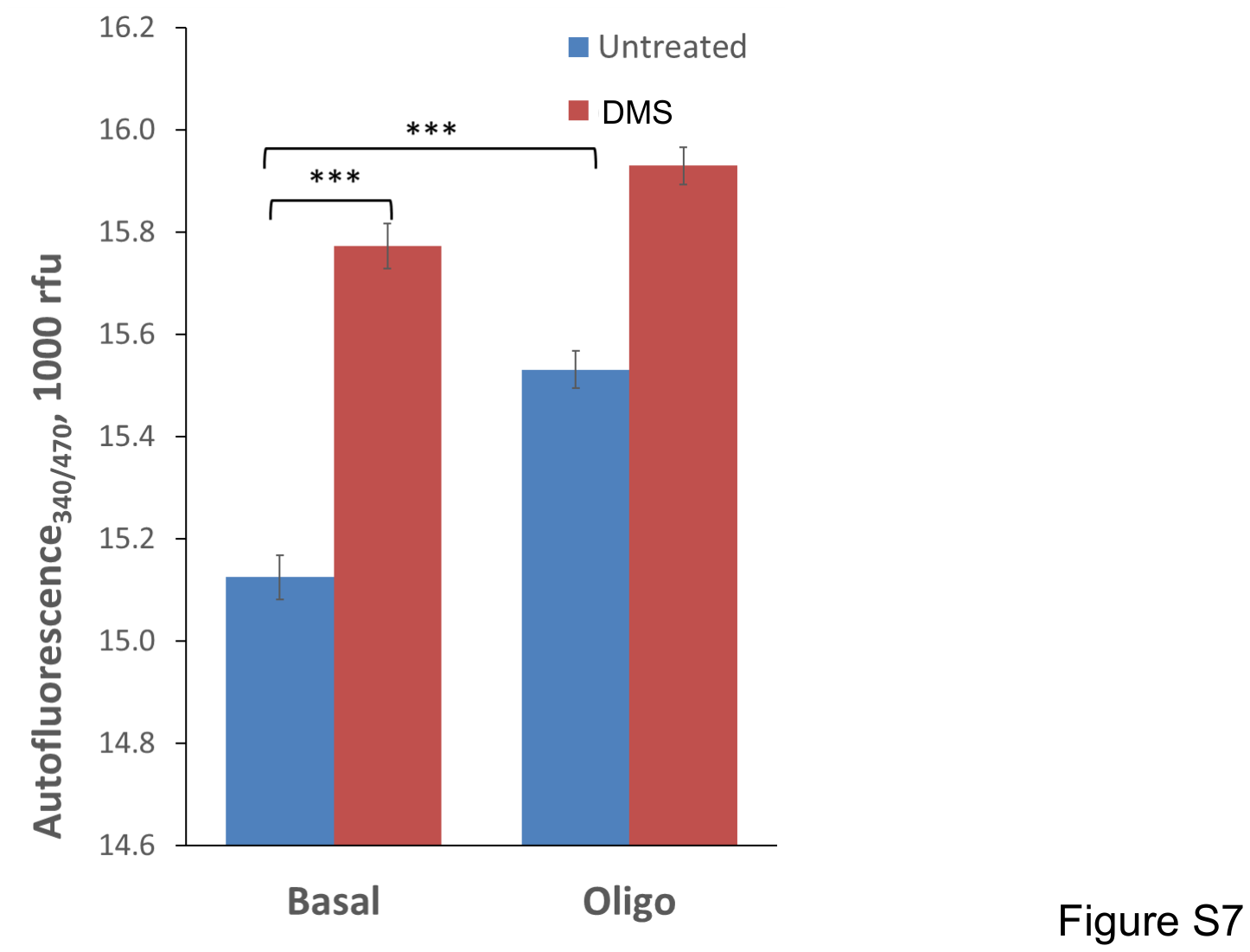

**Figure S7*.* Autofluorescence assay of relative NADH/NAD^+^ ratio in hiN**

Effect of dimethyl succinate (DMS, 5mM) on reductive status of mitochondrial pyridine nucleotides in AD-hiN. Autofluorescence of pyridine nucleotides (yielding the relative NADH/NAD^+^ ratio) was measured as described in METHOD DETAILS. Data are mean and SEM of intensity of individual mitochondrial puncta (n = 200 per field), shown as relative fluorescence units (rfu at 340/470 nm excitation/emission); ***p < 0.001 by Student’s t-test. Additions: Untreated for respiratory basal state; Oligo, 2 µg/ml oligomycin for resting state (State 4).

**Table S1. Data from the organomercury-chemoselective enrichment/MS platform**

**(Tab 1)** EXCEL spreadsheet labeled “Human Brains Information” has Demographics of human AD and control brains analyzed.

**(Tabs 2 and 3)** EXCEL spreadsheets labeled “Control” and “AD” contain SNO-sites found on proteins by MS in Control and AD brains, respectively.

**(Tabs 4 and 5)** EXCEL spreadsheets, labeled “Unique to Control” and Unique to AD” show SNO-sites on proteins found only in Control of AD brains, respectively.

**(Tabs 6-8)** EXCEL spreadsheets labeled “Shared Proteins and Sites,” “Shared Proteins New Sites Control,” and “Shared Proteins New Sites AD” contain lists of proteins with identical S-nitrosylated proteins and SNO-sites, shared S-nitrosylated proteins but with additional SNO-sites found only in Control brains, and shared S-nitrosylated proteins but with additional SNO-sites only in AD brains, respectively.

**(Tab 9)** EXCEL spreadsheet labeled “Venn Data Summary” illlustrates numbers of S-nitrosylated proteins and peptides found in Control human brains, AD human brains, and shared between them.

**(Tab 10)** EXCEL spreadsheet labeled “Shared Sites p < 0.05” shows S-nitrosylated proteins and their SNO-sites that are significantly (p < 0.05) up or downregulated in Control and AD human brains.

**(Tab 11)** EXCEL spreadsheet labeled “TCA Enzymes” contains a list of S-nitrosylated enzymes related to the TCA cycle and their SNO-sites and whether they are differentially S-nitrosylated in AD compared to Control human brains.

**(Tab 12)** EXCEL spreadsheet labeled “GO_BP_CONTROL” shows a list of GO biological process terms (with fold enrichment and false discovery rate [FDR]) that manifest S-nitrosylated proteins from Control human brains.

**(Tab 13)** EXCEL spreadsheet labeled “GO_BP_CONTROL” shows a list of GO biological process terms (with fold enrichment and false discovery rate [FDR]) that manifest S-nitrosylated proteins from AD human brains.
